# Comparison and evaluation of methods to infer gene regulatory networks from multimodal single-cell data

**DOI:** 10.1101/2024.12.20.629764

**Authors:** Pau Badia-i-Mompel, Roger Casals-Franch, Lorna Wessels, Sophia Müller-Dott, Rémi Trimbour, Yunxiao Yang, Ricardo O. Ramirez Flores, Julio Saez-Rodriguez

## Abstract

Cells regulate their functions through gene expression, driven by a complex interplay of transcription factors and other regulatory mechanisms that together can be modeled as gene regulatory networks (GRNs). The emergence of single-cell multi-omics technologies has driven the development of several methods that integrate transcriptomics and chromatin accessibility data to infer GRNs. While these methods provide examples of their utility in discovering new regulatory interactions, a comprehensive benchmark evaluating their mechanistic and predictive properties as well as their ability to recover known interactions is lacking. To address this, we built a comprehensive framework, Gene Regulatory nETwork Analysis (GRETA), available as a Snakemake pipeline, that includes state of the art methods decomposing their different steps in a modular manner. With it, we found that the GRNs were highly sensitive to methods’ choices, such as changes in random seeds, or replacing steps in the inference pipelines, as well as whether they use paired or unpaired multimodal data. Although the obtained networks performed well in predictive evaluation tasks and partially recovered known interactions, they struggled to capture causal relationships from perturbation assays. Our work brings attention to the challenges of inferring GRNs from single-cell omics, offers guidelines, and presents a flexible framework for developing and testing new approaches.

**Graphical Abstract:** 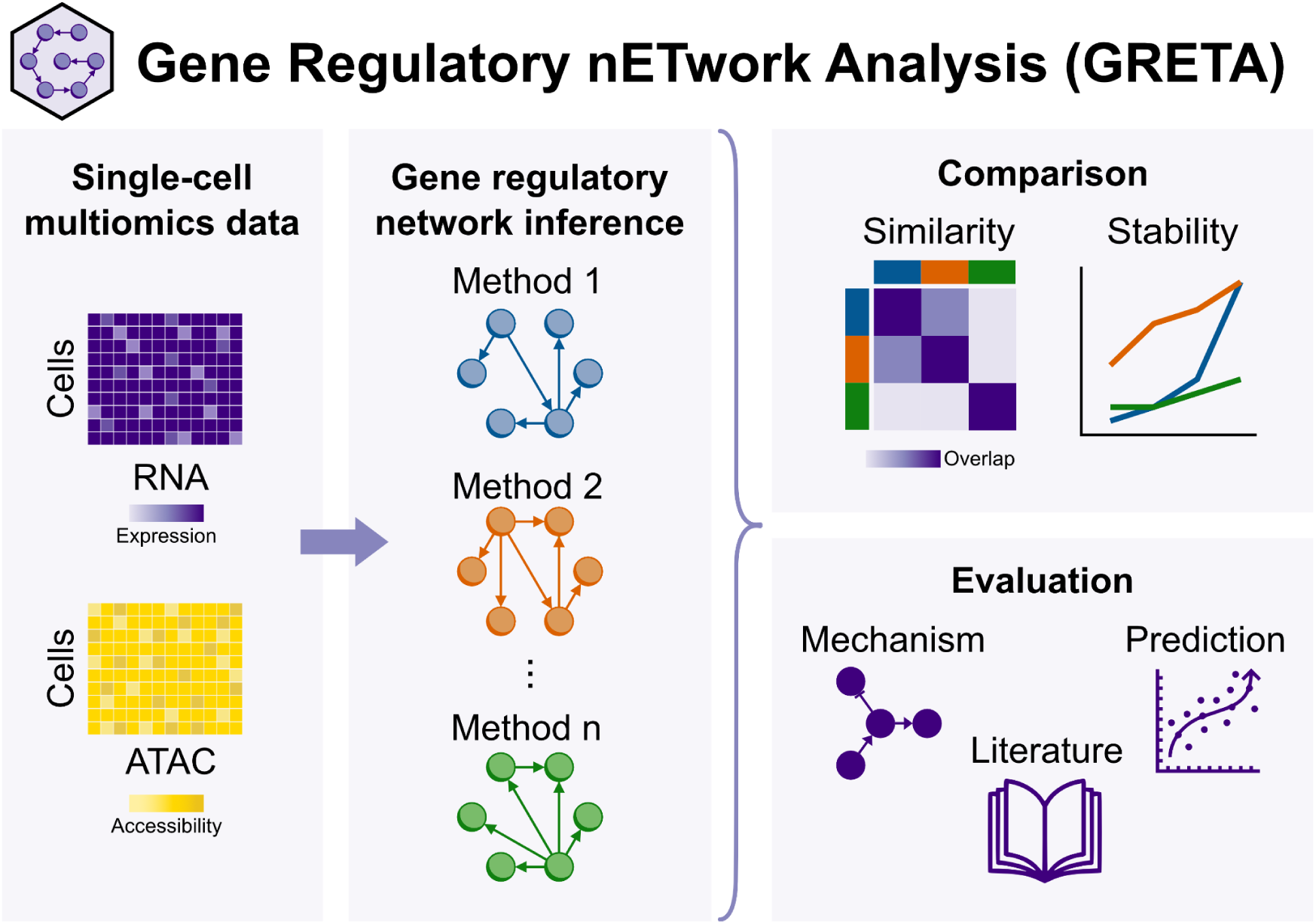

## 1. Introduction

Cells regulate transcription in response to intracellular and extracellular signals, primarily through transcription factors (TFs), a family of proteins that bind DNA to influence gene expression^1^. Genomic DNA is tightly packed into chromatin, rendering most genes inaccessible. Pioneer TFs can initiate changes in DNA accessibility, while other TFs bind to proximal regions like promoters or distal cis-regulatory elements (CREs) to recruit and stabilize the RNA polymerase complex for mRNA synthesis from the gene body^1^. Alternatively, TFs can also reduce gene accessibility, impeding the transcription machinery and suppressing gene expression^1^.

The interactions between DNA regions, TFs, target genes and other molecules form intricate regulatory circuits, which can be modeled as gene regulatory networks (GRNs)^2–4^. These networks are typically represented as graphs, where nodes represent TFs and genes, and edges indicate regulatory relationships between them. These interactions can be positive, where a TF activates expression of a gene, or negative, where it inhibits it. The group of target genes regulated by a TF is known as a TF regulon, and the collection of regulons form a GRN. The analysis of GRNs provides valuable insights into the processes by which cellular identity is established, maintained, and altered in disease^5,6^.

Historically, GRNs have been inferred de-novo from experimental data^4^, particularly bulk omics data, as well as from literature sources^7–9^, which compile findings from multiple assays. The emergence of single-cell multi-omics technologies^10^, particularly the joint profiling of single-cell transcriptomics (snRNA-seq) and chromatin accessibility (Assay for Transposase-Accessible Chromatin using sequencing; snATAC-seq), has driven the creation of advanced computational methods that integrate these multimodal data types. Although they use a variety of strategies, we have identified four shared inference steps that methods use to link TFs to target genes: processing of candidate TFs, CREs and genes; assigning CREs to neighbouring genes; predicting TF binding on CREs; and identifying the final TF-gene interactions through mathematical modelling. Contrary to literature-derived GRNs, these methods potentially enable the inference of highly context-specific regulatory interactions. TF-gene interactions can vary depending on the chromatin state, making these approaches particularly promising for improving GRN reconstruction^11^. This is especially relevant given that transcriptomics-only methods have shown significant limitations in past benchmarks^12,13^.

The joint modeling of these data types also comes with its own challenges, as current profiling technologies tend to yield sparse and noisy readouts, they require computational integration approaches, and have an increased computational cost due to the larger number of features considered^11^. Although multimodal GRN inference involves a sequence of shared steps across methods, their original implementations tend to be rigid, not allowing the use of other approaches in their pipeline. As a result, their application and comparison remain difficult tasks. Additionally, without technologies to directly measure interactions between regulatory elements and their effects on gene expression at large-scale, evaluating the resulting networks remains challenging^11–13^. A comprehensive benchmark of the existing methods is lacking, and it is thus still unclear how well these different methods perform, or even to which extent this joint modeling improves inference compared to methods that only use transcriptomics.

Here, we systematically compared six recently published multimodal single-cell GRN methods^14–19^ alongside four alternative methods^8,9,20^. We first assessed the stability and overlap between methods, followed by an analysis of how paired versus unpaired multiomics data affect their results. Next, we examined the influence of different inference step combinations on GRN inference by modularizing the original inference steps of each multimodal method and testing all possible method-step combinations. Lastly, we developed a comprehensive set of mechanistic, predictive, and knowledge-based metrics to evaluate the original and modularized versions of the methods. All results were generated using Gene Regulatory nETwork Analysis (GRETA, https://github.com/saezlab/greta), a free open-source framework to compare and evaluate multimodal GRN methods, implemented in a modular Snakemake^21^ pipeline that allows to easily run any combination of them. We found a high heterogeneity in the inferred GRNs, that is highly dependent on the choice of inference steps, and modest performance in benchmarks. Overall, our work underscores the complexity of GRN inference, offers guidelines to apply existing methods, and provides a flexible framework for the field to develop and benchmark new approaches.

## 2. Results

### 2.1. GRETA: A framework to systematically build, compare and evaluate gene regulatory networks from single-cell RNA and ATAC data

Multimodal methods that infer gene regulatory networks (GRNs) from single-nuclei transcriptomics (snRNA-seq) and chromatin accessibility (Assay for Transposase-Accessible Chromatin using sequencing, snATAC-seq) employ various strategies, that we have synthetized and classified into four main inference steps: (1) preprocessing data to identify candidate transcription factors (TFs), cis-regulatory elements (CREs), and genes; (2) linking CREs to neighboring genes based on genomic proximity to the transcription starting sites (TSSs) of genes, creating CRE-Gene edges; (3) predicting TF binding to CREs based on motif sequence similarity, forming TF-CRE-Gene triples; and (4) mathematical modeling to derive simplified TF-Gene interactions^11^. To evaluate the agreement between distinct GRN reconstruction methods from multimodal snRNA-seq and snATAC-seq data, and to assess the impact of individual inference steps, we built Gene Regulatory nETwork Analysis (GRETA, https://github.com/saezlab/greta), a free and open-source modular Snakemake pipeline that modularizes GRN inference steps, comparing and evaluating them (**Fig. 1**).

**Figure 1.**
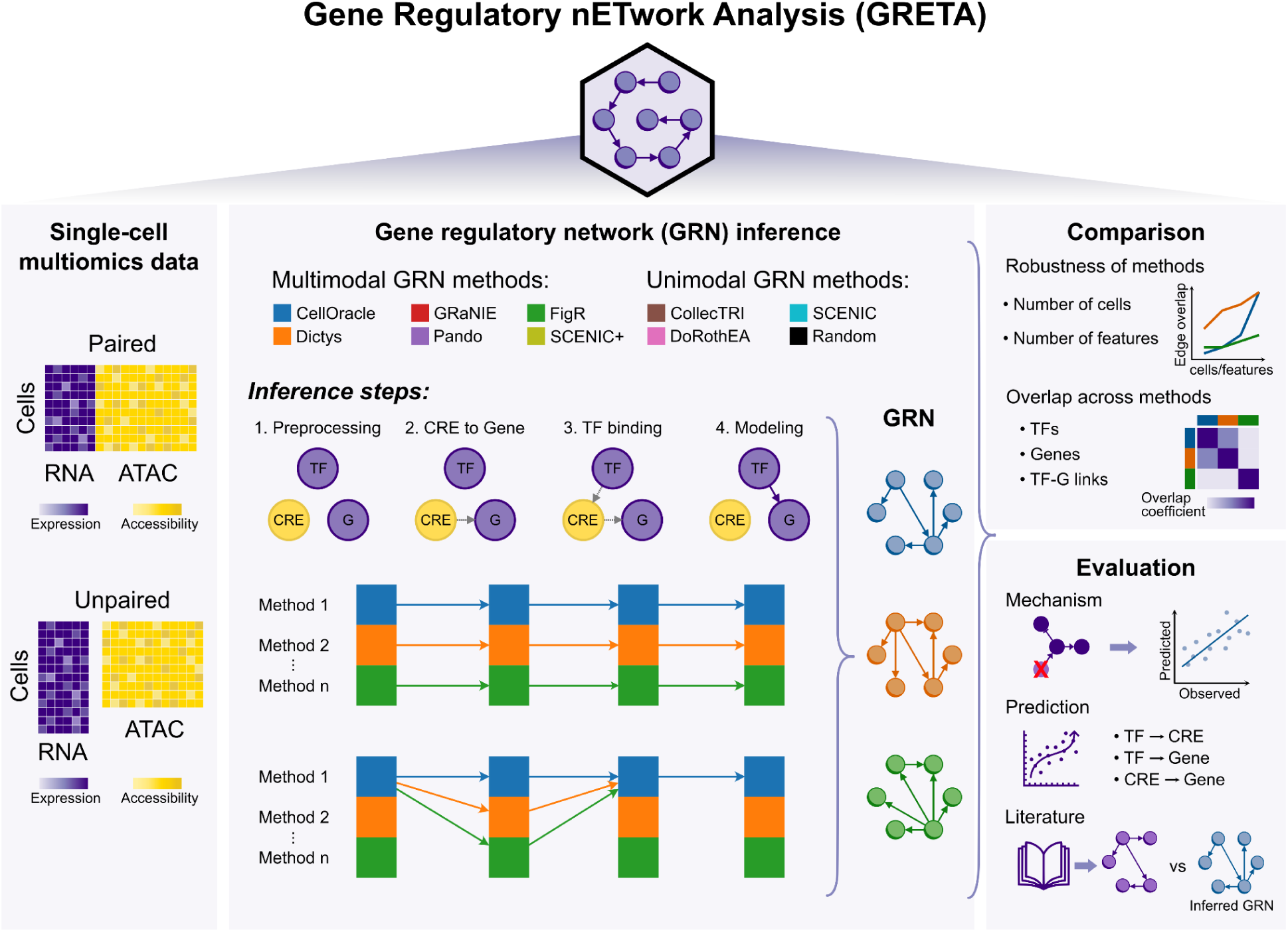
Gene Regulatory nETwork Analysis (GRETA). GRETA is a flexible Snakemake pipeline designed to infer, compare, and evaluate gene regulatory networks (GRNs) from single-cell multimodal RNA-seq and ATAC-seq data. Its modular design allows for any combination of inference steps. GRNs generated with GRETA can be compared across TFs, edges, and target genes to assess stability under different conditions. Additionally, GRETA includes a collection of evaluation metrics categorized into three groups: mechanistic, predictive, and literature-based metrics.

From paired or unpaired multimodal single-cell RNA-seq and ATAC-seq data, GRETA systematically executes the original implementation of six state-of-the-art multimodal methods^11^ (*CellOracle*^14^, *Dictys*^15^, *FigR*^16^, *GRaNIE*^17^, *Pando*^18^, and *SCENIC+*^19^). It also provides their separate inference steps, enabling combinations to generate novel inference approaches. In addition, GRETA executes unimodal methods: *SCENIC*^20^, the state-of-the-art GRN inference method for single-cell transcriptomics; two prior knowledge based GRNs, *CollecTRI*^9^ and *DoRothEA*^8^; and generates random networks with topological properties similar to the ones of inferred GRNs (**Methods**).

GRETA enables systematic comparison of GRNs at the TF, edge, and target gene levels. To account for differences in network sizes, it uses the overlap coefficient, defined as the intersection size between two sets divided by the smaller set’s size. Using this statistic, it can also measure the robustness of methods by sampling distinct random seeds during inference, or by downsampling features or cells (**Methods**).

To extend GRN benchmarking strategies beyond the recovery of TF binding peaks measured by ChIP-seq data and to incorporate the novel multimodal GRNs with distal CREs, GRETA offers a diverse collection of metrics grouped into three categories: mechanistic, predictive, and literature-based (**Fig. 2**). Detailed objectives and evaluation procedures for each metric are provided in **Table 1** and **Methods.** Mechanistic metrics evaluate whether GRNs capture causal relationships from perturbation assays by assessing three aspects: their ability to identify perturbed TFs using enrichment scores from gene signatures (“TF activity”), their accuracy in predicting gene expression changes after perturbation (“Forecasting”), and their capability to identify patterns of TF expression in steady states similar to the ones observed in different cell types (“Steady states”) (**Methods**). Predictive metrics assess how well GRN topology models observational single-cell RNA-seq and ATAC-seq data (“Omics”) and gene set enrichment scores (“Gene sets”) (**Methods**). Literature-derived metrics measure the extent to which GRNs incorporate prior knowledge from databases, focusing on several aspects: inclusion of known TF markers (“Marker TFs”); TF-TF protein-protein interactions based on shared target genes (“TF-TF pairs”); TF binding events to CREs validated by independent ChIP-seq data (“TF binding”); CREs identified as functionally active through biochemical assays, evolutionary conservation, or genetic variant presence; and CREs linked to genes via expression quantitative trait locus (eQTLs) (**Methods**).

**Figure 2.**
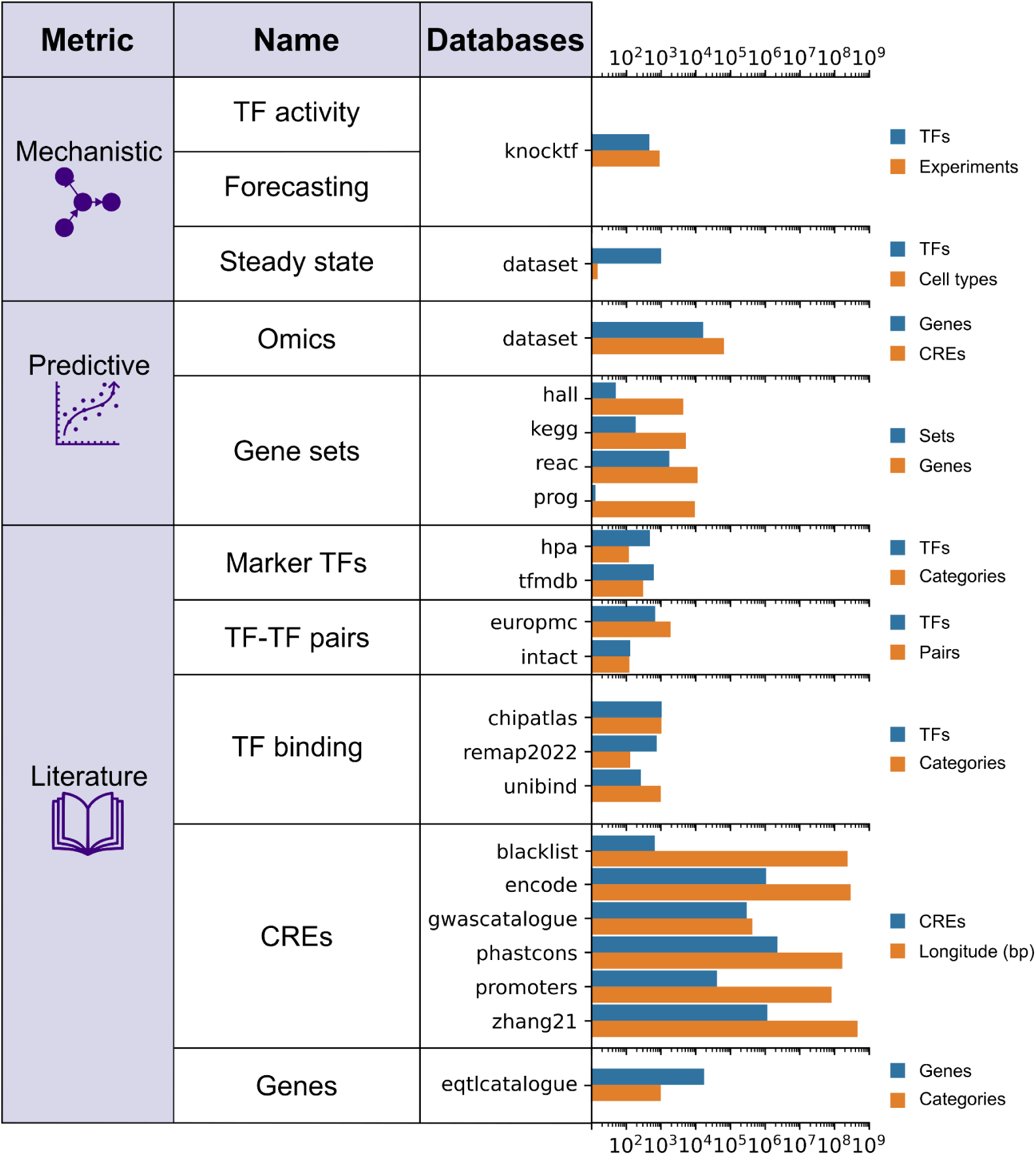
Evaluation metrics and processed databases. List of evaluation metrics, the databases employed to calculate evaluation scores and their statistics such as number of elements, or number of cell-type or tissue labels. The “Steady State” and “Omics” metrics rely on the used single-cell data rather than a database.

After processing all evaluation databases, we observed a high overlap of coverage for target genes, a moderate coverage for TFs, and a low coverage for overlapping genomic regions across resources (mean overlap coefficient genes = 0.7528; mean overlap coefficient TFs = 0.5652; mean overlap coefficient base pairs = 0.2515)(**Extended Data Fig. 1a-c**).

### 2.2. Systematic comparison and stability quantification of GRNs inferred from multiple methods

We used GRETA to reconstruct the GRN of a publicly available, annotated single-cell multiome dataset of human pituitary gland^22^ using the original implementations of the six multimodal GRN reconstruction methods, *SCENIC*, two prior knowledge GRNs, and a random GRN (**Methods**).

First, we evaluated the robustness of the methods in reproducing GRN inference across three different random seed runs using the same pituitary dataset. Most methods consistently recovered the same TFs and target genes through each run (mean overlap coefficient TFs = 0.883; mean overlap coefficient genes = 0.855), as well as their interactions (mean overlap coefficient edges = 0.719)(**Extended Data Fig. 2a**). However, *Dictys*, *SCENIC+*, *SCENIC,* and, as expected, the random network, showed substantial decreases in overlap of edges, with overlap coefficient values around or below 0.5 (**Fig. 3a**). Next, we evaluated the consistency of interaction weights across replicates for the methods that did not achieve a complete overlap (overlap coefficient = 1) across random seeds. Most methods showed strong correlations, reflecting stable interaction inference (median Pearson’s ρ = 0.936) (**Extended Data Fig. 2b**). However, *Dictys* displayed correlation values below 0.5, with highly weighted TF-Gene interactions reversing their direction between runs. Because the runs of the random network yielded no intersection between them, no correlation could be computed. These findings suggest that the probabilistic nature of certain methods can affect GRN inference and impact downstream analyses.

**Figure 3.**
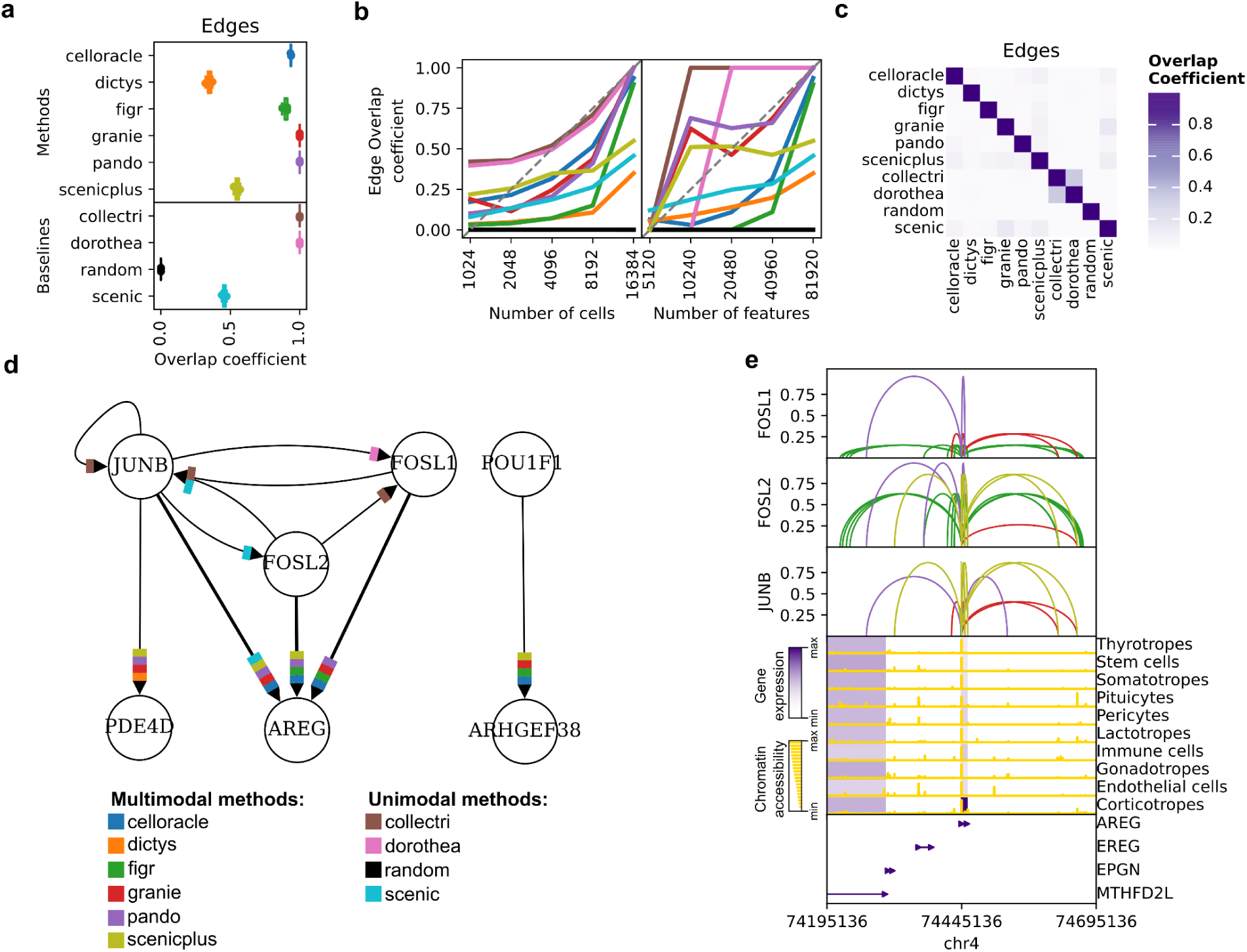
Stability and overlap of multimodal GRN inference methods. **a,** Overlap coefficients of edges between three runs with different random seeds per method. **b,** Overlap coefficients of edges between GRNs inferred from increasingly downsampled datasets at the feature or cell level with the one inferred using the full dataset. **c,** Overlap coefficients of edges across methods. **d,** A shared network across multimodal GRNs with overlapping edges from the unimodal ones. Regulatory interactions to the AREG gene are highlighted by a wider link. **e,** Genomic neighborhood of AREG showing the gene annotations (bottom), the mean gene expression and chromatin accessibility of CREs across cell types (middle), and predicted TF-CRE interactions for AREG’s TFs. Link heights indicate the quantile of the TF-Gene scores inside each method.

We next examined how data size affects GRN reconstruction, since the number of cells or features retained from the same sampled tissue can vary depending on the experimental protocol, and the single-cell data processing framework and its chosen parameters^23^. The pituitary gland dataset was progressively downsampled across four decreasing values in terms of the number of highly variable features or cells, and GRNs were inferred as previously described. For each downsampling level, three different random seeds were used. The resulting networks were compared to those generated from the full dataset at the TF, edge, and target gene levels using the overlap coefficient. A stability score was calculated based on the area under the curve of overlap coefficients, with values close to one indicating minimal impact to downsampling and values close to zero indicating substantial impact (**Methods**). Reducing numbers of cells and features did not drastically impact the recovery of TFs and genes, with a moderate reduction in overlap coefficients (mean stability score TFs = 0.714, genes = 0.59) (**Extended Data Fig. 2c,d**). However, we observed a sharp decline in similarity at the edge level across all methods (mean stability score edges = 0.349) (**Fig. 3b**, **Extended Data Fig. 2d**). Measurements of running times and memory usage during the subsampling strategy underlined the substantial computational resource requirements of multimodal methods (**Supplementary note 1**, **Extended Data Fig. 2e**). These findings show that data coverage strongly impacts network structure, especially interactions, emphasizing the need to optimize preprocessing to retain key network features.

Next, we compared the GRNs obtained from the same dataset with different methods. Most methods inferred GRNs with similar numbers of TFs, edges, and number of target genes per TF (median TFs = 235, edges = 3,010, genes = 1,644, regulon size = 19.56)**(Extended Data Fig. 3a)**. *Dictys* and *SCENIC*, however, produced much larger networks with over 20k edges, nearly 10k target genes, and regulons exceeding 50 genes. Although *SCENIC+* generated a smaller network, its regulon size was comparable to that of *Dictys* or *SCENIC*. Additionally, methods linked genes to CREs that were located at different genomic distances, even though all methods were set to use the same distance parameter (± 250 kb from TSS^1^) (median CRE distance to TSS = 32.37 kb)(**Extended Data Fig. 3b**). In particular, *CellOracle* mostly only recovered promoter regions close to the TSS, while *Pando* and *SCENIC+* sometimes identified distal interactions outside the distance limit. In summary, the GRNs obtained from different methods have substantial topological differences.

When comparing the overlap of the GRNs obtained with different methods, they showed moderate agreement at the TF and target gene levels (median overlap coefficient TFs = 0.536; median overlap coefficient genes = 0.569)(**Extended Data Fig. 3c**), but minimal overlap at the edge level (median overlap coefficient edges = 0.021)(**Fig. 3c**). We also found notable differences between the distinct TSS annotations used across methods (mean genomic overlap coefficient = 0.7532)(**Extended Data Fig. 3d**). Particularly, annotations from *CellOracle*, *FigR*, and *Pando* had the fewest overlaps to the others (mean genomic overlap coefficient *CellOracle* = 0.6389, *FigR* = 0.59, *Pando* = 0.6334). While the rest of methods use similar releases of Ensembl’s hg38 gene annotation version^24^, *CellOracle* uses a custom annotation inferred using Homer^25^, *FigR* uses hg19 RefSeq, and *Pando* uses an old Ensembl release (May of 2017). These results show that, while these methods share similar inference steps, their algorithmic choices and use of prior knowledge influence GRN reconstruction.

Although the methods showed limited overlap at the edge level, for a small regulatory network involving the TFs FOSL1, FOSL2, JUNB, among others, each regulatory interaction was recurrent in at least four out of the six multimodal methods at a time (**Fig. 3d**, **Extended Data Fig. 3e**). GRN methods jointly identified *AREG*, a signaling ligand of the epidermal growth factor receptor^26^, as a highly regulated target gene by these TFs. *AREG* was neither present in the literature-derived GRNs nor in the random network. These results illustrate how GRN inference methods can uncover previously uncharacterized regulatory interactions.

To understand how different methods justified the regulatory links to *AREG* we visualized their predicted TF-CRE-Gene interactions on the genome (**Fig. 3e**). While *AREG*’s promoter region was accessible across all cell types, its expression was restricted to corticotropes, a pituitary gland cell type that secretes stress hormones upon hypothalamic stimulation^27^. This suggested that *AREG*’s expression was influenced by context-specific distal cis-regulatory interactions. Comparing the CREs used by each method to support the same TF-Gene interactions revealed notable differences, illustrating variability in how methods infer regulatory links.

Altogether, the results in this section highlight variability in network size and connectivity across methods, reflecting differences in their inference steps and algorithmic strategies.

### 2.3. Evaluating the impact of paired vs unpaired multiome data in GRN inference

Another key element to consider when performing GRN inference is the nature of the multimodal data being used. Whether the data are truly paired, measuring both, snRNA-seq and snATAC-seq from the same cells, or unpaired, measuring them from different cells, has the potential to impact GRN reconstruction^28^. While some methods, such as *CellOracle* and *GRaNIE*, are designed to handle both paired and unpaired data, others are limited to paired data and rely on computational strategies to integrate the modalities when working with unpaired datasets. To evaluate the impact of these differences on GRN inference, we again used the human pituitary gland dataset, which uniquely includes profiles from the same donor generated with both paired and unpaired multiome technologies, making it particularly suitable for this comparison **(Fig. 4a).**

**Figure 4.**
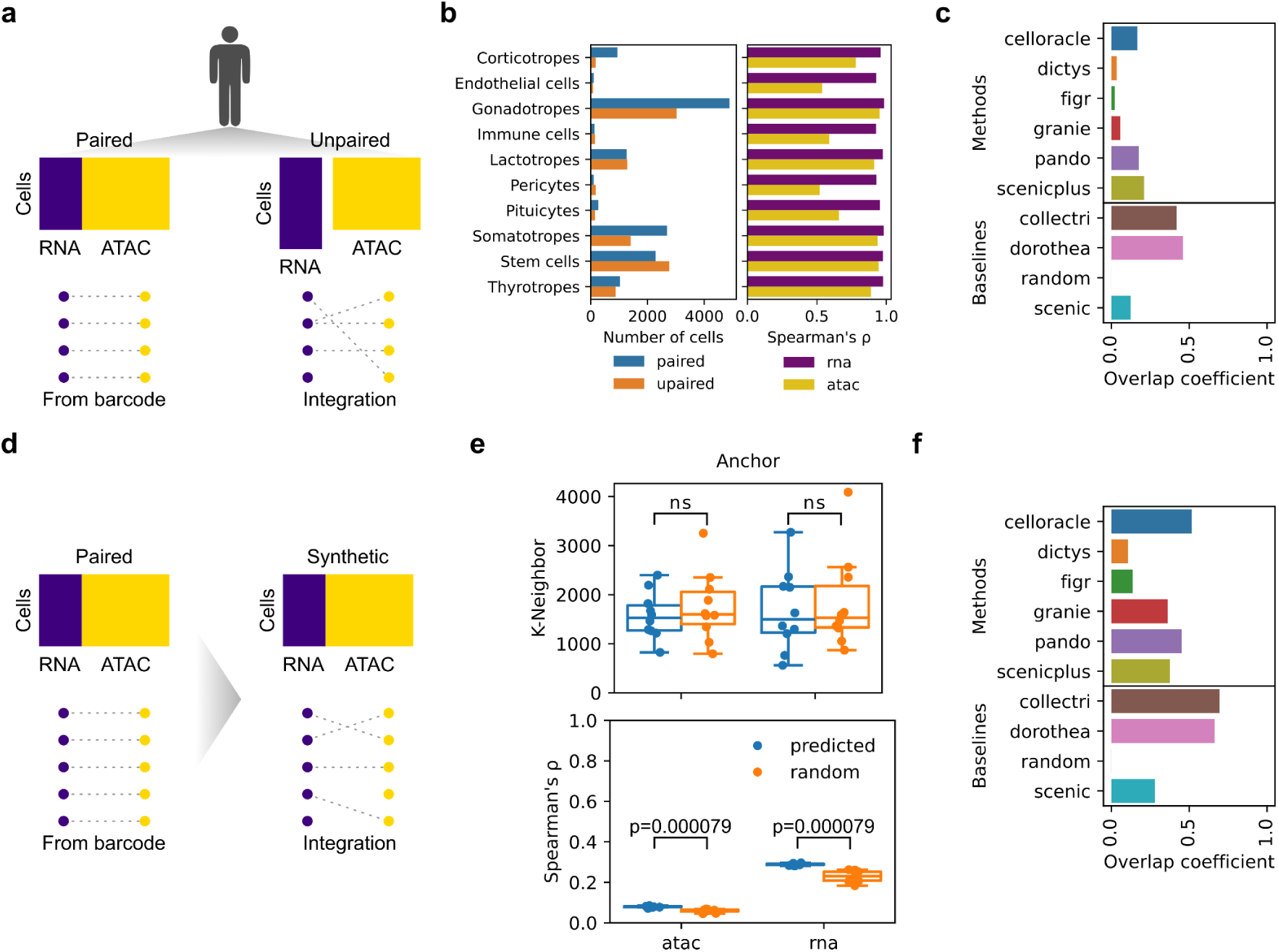
Impact of paired vs unpaired multimodal data in GRN inference. **a**, Paired and unpaired single-cell multiomics were measured for the same pituitary gland. In paired data, the mapping between modalities cells is known, while in unpaired the number of cells can be different and requires integration between modalities. Missing or non-straight lines denote imperfect matching. **b,** Number of cells per cell type for both datasets (left), Spearman’s ρ between the molecular readouts of the paired and unpaired datasets per cell type (right). **c,** Overlap coefficient of edges between the GRNs inferred from the paired and unpaired dataset. **d,** By treating the paired pituitary dataset as if it were unpaired, a synthetic integrated dataset was created by matching cells using FigR’s integration approach. Missing or non-straight lines denote imperfect matching. **e,** Mean k-neighbour for each cell type of the predicted and random cells from the same cell type (top), and the mean Spearman ρ between the predicted and random cells across data modalities. Anchor refers to which modality was used in the obtained mapping to make the comparisons. **d,** Overlap coefficient of edges between the GRNs inferred from the paired and synthetic dataset.

To integrate the unpaired dataset, we used *FigR*’s integration approach based on optimal transport, as it is one of the few methods that return a one-to-one mapping between barcodes (**Methods**). After identical processing, we observed that the paired and unpaired datasets were similar (**Extended Data Fig. 4a**), both at the level of cell type abundances (cosine similarity = 0.94) and molecular readouts (RNA mean Spearman ρ = 0.9586; RNA mean FDR < 2.2×10^−16^; ATAC mean Spearman ρ = 0.7722; ATAC mean FDR < 2.2×10^−16^)(**Fig. 4b**). We also noticed that both runs had a similar sequencing depth and total number of features (mean log1p total counts = 8.01; mean log1p number of genes = 7.36)(**Extended Data Fig. 4b**). However, we found that the unpaired dataset showed a higher mean sequencing depth (two-sided Rank sums test, N per group= 10 cell types; W = 2.57; BH FDR = 0.01) and mean number of features (two-sided Rank sums test, N per group= 10 cell types; W = 2.87; BH FDR = 0.0162) across cell types in snRNA-seq. We then compared the GRNs inferred from both datasets for each method, and we found low overlap between the edges of the networks (mean overlap coefficient edges = 0.1471)(**Fig. 4c**). These results suggested that, while paired and unpaired multiome datasets retain global cell-type properties, the computational integration of unpaired data introduces differences that affect the inferred GRNs.

To assess the effects of data integration on GRN inference, we compared the original paired multiome pituitary dataset with a synthetic paired dataset, generated by treating the original data as unpaired and integrating it using *FigR’s* pairing method (**Methods, Fig. 4d**). Synthetic pairing correctly assigned only 87 cells to their corresponding original cell from the paired information (0.63 % of total cells). Although integration generally paired cells across modalities within the same cell type **(Extended Data Fig. 4c)**, the assigned pairs were as distant from the original cell as randomly assigned pairs within the same cell type both at the RNA-seq (one-sided Rank sum test; median k-neighbor of predicted cells in RNA per cell type = 1,495.71; median k-neighbors of random cells in RNA per cell type = 1,530.25; N per group = 10 cell types; W = -0.378; P = 0.3527) and ATAC-seq (one-sided Rank sum test; median of k-neighbor predicted cells in ATAC per cell type = 1,527.31; median k-neighbor of random cells in ATAC per cell type = 1,597.78, N per group = 10 cell types; W = -0.453; P = 0.3251) (**Fig. 4e**) level. Nevertheless, synthetically paired cells showed a higher correlation of RNA and ATAC features compared to randomly assigned pairs within the same cell-type (one-sided Rank sum test; median Spearman’s ρ of predicted cells in RNA per cell type = 0.2874; median Spearman’s ρ of random cells in RNA per cell type = 0.23; N per group = 10 cell types; W = 3.7796; P = 7.85e-05)(one-sided Rank sum test; median Spearman’s ρ of predicted cells in ATAC per cell type = 0.0795; median Spearman’s ρ of random cells in ATAC per cell type = 0.0619; N per group = 10 cell types; W = 3.7796; P = 7.85e-05)(**Fig. 4e**). Comparison of GRNs from the paired and synthetic paired datasets revealed low overlap of interactions (mean overlap coefficient edges = 0.35; **Fig. 4f)**. These findings suggested that while integration accurately groups cells by cell type, it fails to align them within finer cell states, reducing the effectiveness of GRN methods that rely on paired multimodal data.

### 2.4. Decomposing multimodal GRN inference

To evaluate the impact of each inference step of multimodal methods in the reconstruction of GRNs, we used a paired multiome dataset of peripheral blood mononuclear cells (**Extended Data Fig. 5a-c**) and systematically replaced individual steps in each original method with those from other methods (**Methods**). To validate our implementations, we compared GRETA’s modularized version of multimodal GRN methods to the originally fixed ones. All methods demonstrated high consistency, with substantial edge overlap coefficients and Spearman’s ρ between the original and GRETA’s implementations (mean edge overlap coefficient = 0.93; mean Spearman’s ρ = 0.989) (**Extended Data Fig. 5d**).

We next combined the inference steps across methods and observed that replacing individual steps with those from other methods had varying impacts on GRN reconstruction. Across different combinations, overlap coefficients for TFs and genes included in the GRNs remained consistently above 0.5, although gene-level overlap showed greater variability (TF mean overlap coefficient = 0.846; gene mean overlap coefficient = 0.8215; **Extended Data Fig. 5e-h**). However, at the edge level, modifying a single analysis step substantially altered the GRN structure, resulting in lower overlap coefficients (mean overlap coefficient = 0.3346; **Fig. 5a**).

**Figure 5.**
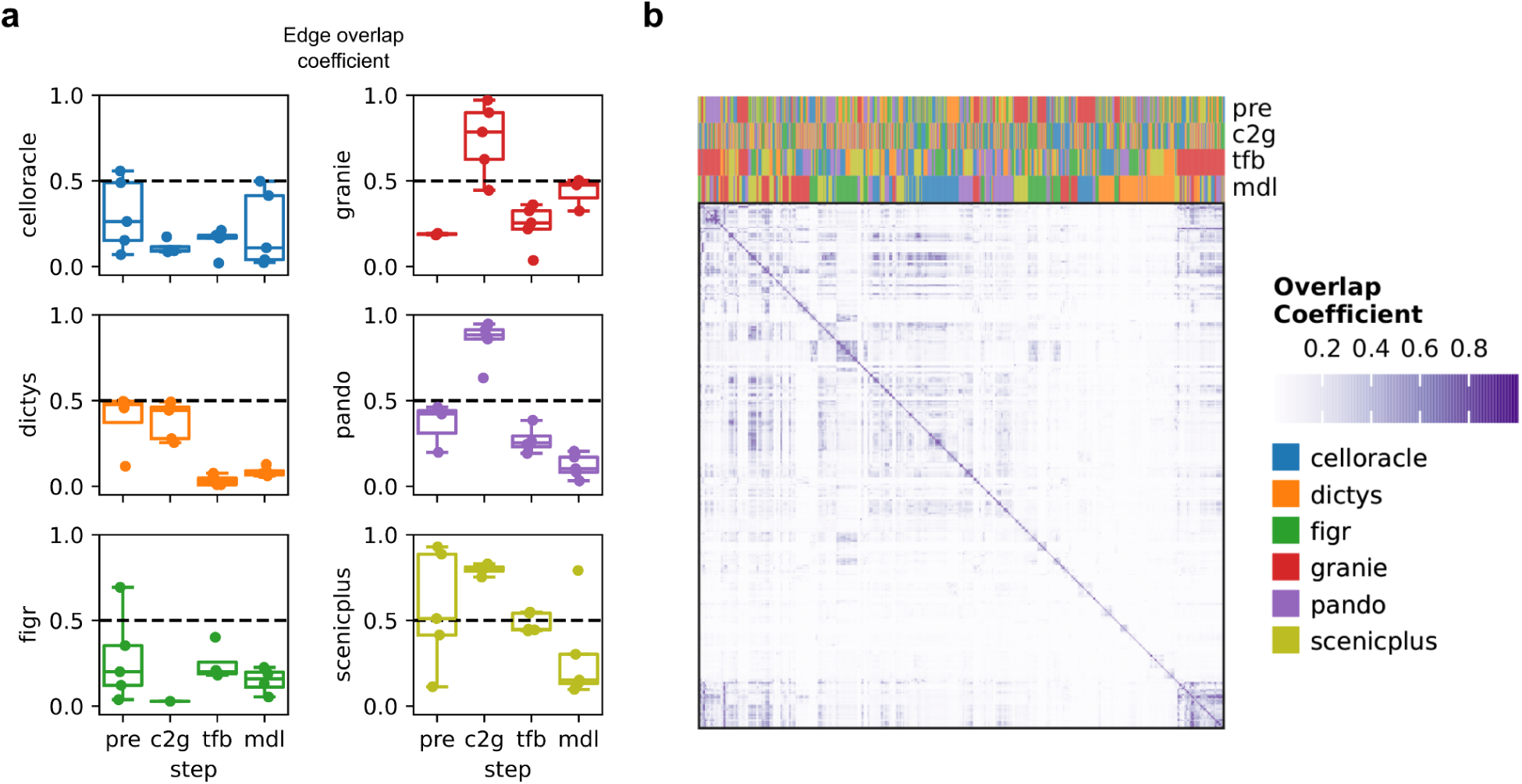
Multimodal GRN inference decomposition. **a,** Effect of changing one of the inference steps from a fixed method pipeline to another method, measured by the overlap coefficient at the edge level. **b,** Pairwise overlap coefficient at the edge level between all possible inference runs combinations. Preprocessing (pre), CRE to gene (c2g), TF binding (tfb) and modeling (mdl).

We next examined whether different runs would converge to a limited set of solutions. To assess this, we performed all possible runs (4^N^, where N was the number of multimodal GRN methods) and clustered them based on pairwise overlap coefficients at the TF, edge, and gene levels. Based on the overlap of TFs and genes used in the GRN, methods clustered according to the TF binding and CRE-to-gene steps, respectively (mean TF overlap coefficient = 0.6532; mean gene overlap coefficient = 0.6171)(**Extended Data Fig. 5**). However, when comparing the similarity of GRN interactions across different inference step combinations, edge-level overlap coefficients were only marginally higher than those observed for random GRNs (mean non-random overlap = 0.0884; mean random overlap = 0.0028; **Fig. 5b**). This suggests that although methods incorporate similar TFs and genes, each combination of them generates a unique solution to the inference problem.

### 2.5. Evaluation of multimodal GRNs

To assess the performance of combinations of steps and original multimodal methods, we used the evaluation metrics defined in **Fig. 2** and **Table 1**, applied to the different runs obtained in **Fig. 5**. For each metric and GRN, precision and recall were calculated based on the metric-specific definitions of true positives (TPs), false positives (FPs), and false negatives (FNs)(**Table 1**). These values were summarized into an F score, with precision weighted ten times more than recall (β = 0.1), denoted F_0.1_. This weighting was chosen because GRN inference methods typically return sparse networks that prioritize precision, as they aim to capture the most confident interactions rather than all possible ones in the ground truth^15^. Given that the ground truth is not perfect, precision becomes more important than recall to ensure the predictions accurately reflect the true regulatory interactions. A weighted Kolmogorov-Smirnov-like statistical test was then used to identify method steps that ranked significantly higher based on the F_0.1_ score (**Methods,** FDR < 0.01)(**Fig. 6a**).

**Figure 6.**
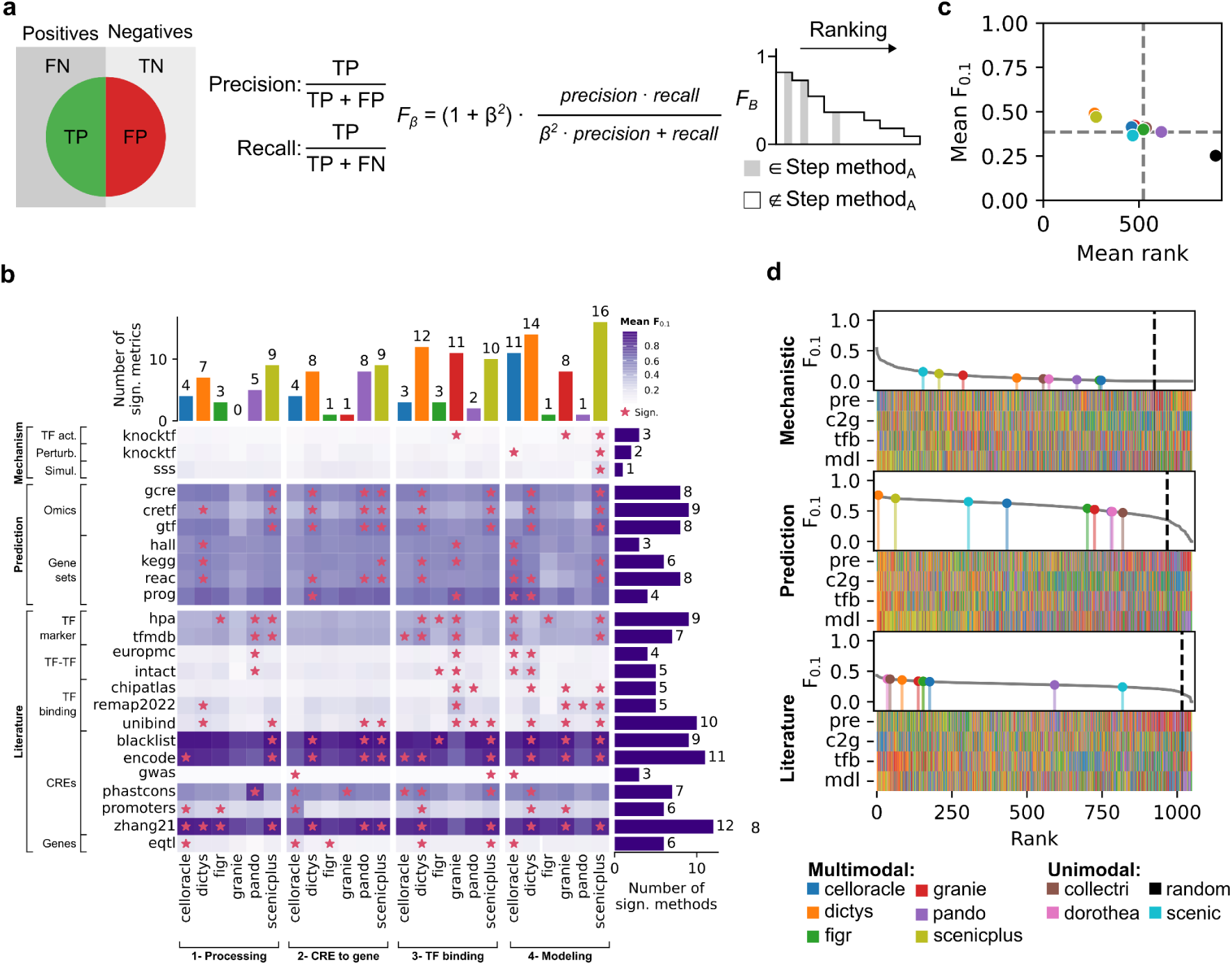
Multimodal GRN evaluation. **a,** Strategy used to assess the performance of each GRN inference step across metrics. Precision and recall were first computed from true positives (TPs), false positives (FPs), and false negatives (FNs). These were then summarized into an F_β_ score, with precision weighted ten times more than recall (β = 0.1). A weighted Kolmogorov-Smirnov-like statistical test was used to assess whether groups of runs sharing a specific method’s inference step were ranked significantly higher than others. **b,** Mean F_0.1_ scores across evaluation metrics and inference steps per method. Asterisks indicate that runs using a method’s step ranked significantly higher than the rest for a particular metric. Barplots indicate the number of significant method-metric pairs grouped per method’s step (top) or per metric (right). **c,** Averaged F_0.1_ scores and ranks across all metrics for the fixed methods plus the unimodal ones. Dashed lines indicate the mean of all possible runs for F_0.1_ and rank. **d,** Averaged F_0.1_ scores based on the three metric categories across runs. Lollipop plots indicate fixed or unimodal pipelines, and color indicates the different methods. The dashed black line indicates the random GRN.

Most GRNs had a moderate performance at recapitulating findings of literature (mean F_0.1_ score = 0.362), including TF markers, TF-TF interactions via shared targets, validated TF binding events, functional CREs identified through assays or conservation, and CRE-gene links supported by eQTLs (**Table 1**). Among the methods evaluated, specific approaches performed better in distinct tasks (**Extended Data Fig. 6**). For recovering marker TFs, *SCENIC+*’s modeling strategy performed better in the HPA database (mean F_0.1_ score = 0.46), while *Dictys*’ TF binding step performed better in the TF-Marker database (mean F_0.1_ score = 0.528). The modeling strategy of *Dictys* yielded the higher scores for both TF-TF databases (mean F_0.1_ score: EuropePMC = 0.2384, IntAct = 0.3059). *GRaNIE*’s TF binding approach outperformed in two of the three TF binding databases (mean F_0.1_ score: ChIP-Atlas = 0.2253, ReMap2022 = 0.3614), while *Dictys*’ preprocessing performed better in the remaining one (mean F_0.1_ score: UniBind = 0.2172). For CRE-related tasks, *CellOracle*’s preprocessing performed best in ENCODE and Zhang21 (mean F_0.1_ score: ENCODE = 0.8678, Zhang21 = 0.8935). *CellOracle*’s CRE to gene strategy also scored highest for recovering GWAS hits, promoter regions and CRE-gene links from eQTL catalog (mean F_0.1_ score: GWAS catalog = 0.00076, promoters = 0.5491, eQTL catalog = 0.2133). Lastly*, Pando*’s preprocessing strategy performed best for evolutionary conservation (mean F_0.1_ score = 0.8295), while *Dictys*’ modeling performed better in recovering CREs outside blacklisted genomic regions (mean F_0.1_ score = 0.9351).

Similarly, most methods performed decently on prediction tasks (mean F_0.1_ score = 0.561), where the objective is to evaluate the prediction of gene expression, regulatory elements accessibility, or biological processes using the network topology (**Table 1**). *SCENIC+*’ modeling strategy achieved the highest scores across all omics tasks (mean F_0.1_ score: Gene ∼ TFs = 0.7057, CRE ∼ TFs = 0.6176, Gene ∼ CREs = 0.7057), whereas *CellOracle*’s modeling strategy outperformed others in all gene set databases (mean F_0.1_ score Hallmarks = 0.6166, Kegg = 0.651, Reactome = 0.671, PROGENy = 0.7627) (**Extended Data Fig. 6**).

All combinations of methods performed poorly in mechanistic tasks (mean F_0.1_ score = 0.0735), aimed at predicting the effects of perturbing TFs or simulating similar cell types (**Table 1**). Within this limited performance, *SCENIC+’* modeling approach outperformed the rest of the methods in all mechanistic metrics (mean F_0.1_ score: TF activity = 0.0818, Forecasting = 0.1341, Steady state = 0.251) (**Extended Data Fig. 6**).

When evaluating the performance of methods at each inference step, the preprocessing strategy of *Dictys* and *SCENIC+*, the CRE-to-gene strategy of *Dictys*, *Pando* and *SCENIC+*, the TF binding strategy of *Dictys*, *GRaNIE* and *SCENIC+*, and the modeling strategy of *CellOracle, Dictys* and *SCENIC+* achieved consistently higher F_0.1_ scores across more tasks compared to other strategies (**Fig. 6b**).

Next, we compared the ranks and F_0.1_ scores of the fixed original multimodal pipelines and the unimodal methods against all possible combinations of runs across the evaluation metrics. We found that the fixed original pipelines for *CellOracle*, *Dictys*, *FigR*, *GRaNIE*, and *SCENIC+* outperformed the mean rank and F_0.1_ score of all method combinations (mean across all combinations: rank < 523.43, F_0.1_ score > 0.3853) (**Fig. 6c**). Notably, *Dictys* and *SCENIC+* outperformed the other fixed methods, as they had the lowest mean ranks (mean rank: *Dictys* = 258.58, *SCENIC+* = 275.91) and higher mean F_0.1_ scores (mean F_0.1_: *Dictys* = 0.49, *SCENIC+* = 0.47). In contrast, *Pando* and all unimodal methods had higher ranks and lower F_0.1_ scores compared to all combinations.

We then evaluated the performance of the fixed runs across the three metric types (**Fig. 6d**, **Extended Data Fig. 7a**). *SCENIC* and *SCENIC+* achieved the lowest ranks and highest F_0.1_ scores in mechanistic metrics (mean ranks: *SCENIC =* 154, *SCENIC+ =* 207; mean F_0.1_ scores: *SCENIC* = 0.1547, *SCENIC+* = 0.1243). For predictive metrics, *Dictys* and *SCENIC+* outperformed others (mean ranks: *Dictys =* 5, *SCENIC+ =* 62; mean F_0.1_ scores: *Dictys* = 0.7594, *SCENIC+* = 0.705). Lastly, *CollecTRI* and *DoRothEA* excelled in literature-based metrics, as expected given that these resources rely on literature, followed by *Dictys* (mean ranks: *CollecTRI =* 44, *DoRothEA =* 35, *Dictys =* 84; mean F_0.1_ scores: *CollecTRI* = 0.371, *DoRothEA* = 0.373, *Dictys =* 0.357).

Given that *Dictys* and *SCENIC+* both produced distinct networks across different runs (**Extended Data Fig. 2b**), with *Dictys* even obtaining anticorrelated results, we evaluated their stability in performance by running them with five different random seeds (**Extended Data Fig. 7b**). The results showed that the obtained F_0.1_ scores were consistent (mean standard deviation: *Dictys* = 0.033, *SCENIC+* = 0.016), suggesting that both methods may identify multiple optimal local minima.

In summary, we found substantial differences between inferred networks. While some approaches performed better than others, overall performance was limited based on our benchmarking criteria.

## 3. Discussion

The accumulation of single-cell datasets profiling both transcriptomics (snRNA-seq) and chromatin accessibility (snATAC-seq) has spurred the development of several methods that integrate these data to infer gene regulatory networks (GRNs). These networks link transcription factors (TFs) to accessible cis-regulatory elements (CREs) that regulate neighboring genes. To systematically compare and evaluate these methods, we built GRETA, a framework that integrates and modularizes the inference steps from multiple multimodal GRN inference methods, as well as some unimodal ones. Using GRETA, we explored the stability and agreement among these methods and their inference steps, their sensitivity to the type of multimodal data used during inference, and their performance across different tasks.

Our results show that these methods have limited robustness, yield substantially different outcomes from one another, and are affected by the type of multimodal data being used. Moreover, even within the limited shared regulatory interactions, we observed differences in which cis-regulatory interactions methods use to make their TF-gene predictions using the same data. These differences likely stem from the distinct empirical decisions made during GRN inference, as well as variation in the prior knowledge resources and genome annotations used by each method. In addition, the high variability across networks reinforces the idea that de novo GRN inference from observational data alone may be an ill-posed problem lacking a uniquely defined solution. Such limitations in current methods complicate the evaluation of novel interactions, as technical variability outweighs biological signals.

Multimodal GRN inference method implementations are complex, often requiring diverse input data formats and challenging installations, with advanced coding skills needed to use them effectively, limiting their accessibility to a broader community. GRETA reduces these barriers by offering a modular pipeline that simplifies workflows, supports reproducibility, and facilitates systematic comparisons and novel combinations of GRN inference steps.

Our analysis of the pituitary paired and unpaired multimodal datasets revealed that, although the two datasets were similar at the cell abundance and molecular levels, the inferred GRNs were substantially different. Single-cell multiomics integration remains a challenging problem that has not yet been fully solved^28^. For instance, in RNA and ATAC integration, CREs are transformed into genes by computing gene scores based on gene promoter accessibility^29^, which are then used to match the two modalities at the gene level. While this strategy may successfully map cells to their matching cell type, promoter accessibility does not necessarily guarantee gene expression^1^, making the mapping of cells to specific cell states a difficult task. Specifically, we observed that cells from the same cell type already exhibit low correlation with each other, suggesting that fine mapping of 1-to-1 cells might be unreachable. Consequently, paired data should be employed whenever possible to improve the accuracy of cell state mapping and GRN inference.

As expected, we found that the inferred GRN topologies can reasonably predict basal gene expression, as they use this information during inference. In contrast, they can only partially capture the regulation of biological processes and prior evidence from the literature. Except for *Pando*, all multimodal methods, on average, outperformed unimodal ones, with *Dictys* and *SCENIC+* consistently performing the best. However, all methods performed poorly in mechanistic tasks, where the objective is to predict perturbations relying on the potential causal structure of the GRNs. This lack of causal properties raises the question of which other modalities or technologies we need for reliable trans-regulation predictions. Estimating causal relationships observed in perturbation data from GRNs built from observational data is intrinsically a challenging problem^32^. This difficulty is compounded with the fact that the process of trans-regulation involves multiple steps that are not fully captured by gene expression or chromatin accessibility data alone. For a TF to regulate a target gene, its transcript must first exit the nucleus via nuclear pores, be translated into protein, undergo activation through post-translational modifications, re-enter the nucleus, interact with other nuclear TFs, and interact with accessible chromatin and favorable conditions at the target transcription start site (TSS) to exert regulatory effects. This complex chain of events is further complicated by processes we may not yet fully understand or have not yet discovered. However, the poor predictive performance of perturbations has also been noted in other evaluations, highlighting similar challenges for both classic regression models^30^ and the latest deep-learning and foundation models^31^. Promising experimental approaches to improve GRN inference include systematic perturbation at scale^33^, TF dosage dynamics^34^, or joint spatial 3D accessibility of chromatin with protein staining^35^. From the modeling side, sequence-based models that predict genomic molecular readouts such as chromatin accessibility could be incorporated into GRN inference methods^36^, as they can refine the TF binding step to be context specific.

This study is, to the best of our knowledge, the most comprehensive attempt to systematically evaluate multimodal GRN inference methods. Benchmarking is a challenging task due to the elusive nature of ground truth^11,37^. Here, we addressed this by combining different indirect evaluation criteria based on different assumptions about gene regulation. Another limitation is that all metrics, except for the predictive ones and “Steady States,” depend on context-specific databases. This limitation prevented us from evaluating the pituitary dataset, as few databases had entries for this organ. Additionally, the results are limited to the methods and datasets we used. We used only human data but, given the availability of extensive data for model organisms such as mouse^38^ or fly^39^, it could be expanded to evaluate the generalizability of our findings to these organisms.

In summary, we present GRETA (https://github.com/saezlab/greta), a framework to facilitate the inference, comparison and benchmarking of GRNs from single-cell multiomics data. Our analyses with GRETA help understand the strengths and weaknesses of existing methods, highlighting the need to interpret results of GRN methods with caution and to reassess the limits of modeling gene regulation from observations of gene expression and chromatin accessibility data. As new data modalities and concepts are developed, we envision GRETA as a collaborative platform that the community uses and contributes to, to advance together the GRN inference field.

## 4. Methods

### snMultiome data processing

Raw transcript count matrices, sample-specific chromatin accessibility fragment files and global barcode annotations were retrieved for each dataset from GEO, CELLxGENE and 10x Genomics portal. Fragment files were processed to assign the corresponding sample names to barcodes, ensuring that they were unique. Each dataset was processed independently using the same processing pipeline described below.

For each sample in a dataset, fragment files were processed into tile matrices using SnapATAC2^40^. Barcodes not present in each dataset annotation metadata were removed, guaranteeing that only good quality barcodes were kept. Tile matrices were concatenated and dataset-specific consensus accessibility peaks (Cis-Regulatory Elements, CREs) were called using SnapATAC2’s implementation of MACS3^41^.

Genes expressed in few cells were removed from transcript count matrices, and cells containing few expressed genes, and (number of genes > 100, number of cells > 3). Gene symbols from transcript count matrices were removed if not present in Ensembl 111^24^. For each dataset, if a gene symbol appeared more than once, the feature with counts detected in the highest number of cells was kept. Barcodes were only kept if they were present in the corresponding dataset annotation. Then, transcript and accessibility matrices were paired into a MuData object^42^.

Genes or CREs with low detection were removed (number of counts detected per barcode ≤ 3). For each omic, counts were normalized by barcode-specific total counts (target sum = 1e4) and log1p transformed with scanpy^43^. Feature selection was performed by calculating highly-variable features for each sample and then keeping the top number of features that were found to be variable in the maximum number of samples (number genes = 2^14^, number CREs = 4 x 2^14^).

Simultaneous dimension reduction using Laplacian Eigenmaps was computed with SnapATAC2. For multi-sample datasets, the obtained spectral embeddings were integrated across samples using harmony-py until convergence^44^. With the corrected embeddings, a neighborhood graph of barcodes was inferred, followed by UMAP dimensionality reduction for visualization purposes. Original raw integer counts for genes and CREs were kept in the layers of the MuData object for downstream analyses.

### snRNA-seq and snATAC-seq pairing

To integrate the unpaired snRNA-seq and snATAC-seq pituitary datasets, we first generated a shared co-embedding using Seurat^45^. For the RNA data, we performed log normalization, selected highly variable genes, and scaled the data. For the ATAC data, we applied term frequency-inverse document frequency normalization, selected highly variable CREs, and scaled them as well. Gene scores were inferred from ATAC data and subsequently processed as RNA. Canonical correlation analysis was then used to derive the shared embedding between RNA and inferred gene scores. Individual cells were paired in a one-to-one manner using *FigR*’s integration approach based on optimal transport. Finally, the matched barcodes were merged into a paired MuData object. The same procedure was used to generate the synthetic paired dataset from the real paired pituitary dataset.

### Original multimodal GRN inference methods implementation

This section explains how each multimodal GRN inference method was applied to each dataset, following the original developers’ recommendations as closely as possible.

Following the documentation of *CellOracle* (v0.16.0)^14^, Cicero^46^ was used to identify CRE-CRE co-accessibility interaction scores with a window size around the transcription starting site (TSS) of 500 kb. CREs were annotated to genes if they were located at its TSS, using a custom annotation generated with Homer^47^. Pairs of CRE-TSS interactions were removed according to their interaction score (score < 0.2). For the remaining CREs, TF binding predictions were computed with the motif matcher algorithm GimmeMotifs^48^ and the motif database CIS-BP^49^ (FPR = 0.02, background length = 200 bp, -log_10_(FDR) > 10). CRE-gene and TF binding results were merged to obtain a scaffold TF-CRE-Gene network. K-Nearest Neighbours (KNN) was performed on the PCs of the gene expression matrix to reduce its sparsity (k = 20). Finally, a single tissue-level GRN was inferred with a Bagging Ridge linear model (alpha = 10, number of top significant edges = 2k).

*Dictys* (v1.1.0)^15^ started by performing a basic quality control filtering (minimum total read count per gene > 50; minimum number of cells expression each gene > 10; minimum total read counts per cell > 200; minimum number of non-zero genes per cell > 100). Then, CRE were linked to neighboring genes based on genomic distance using Ensembl’s (release 107) gene annotation^24^ (window size = 500k). Wellington^50^ and Homer^47^ were then applied to identify TF footprints and enriched TF binding sites from HOCOMOCO v11^51^ within the accessibility profiles of each pre-defined cluster. TFs were assigned to CREs by aggregating results from both methods, and the outcomes were averaged across clusters. The CRE-gene and TF-CRE associations were merged to obtain a scaffold TF-CRE-Gene network. Finally, a GRN was inferred by predicting gene expression using stochastic process models, retaining only interactions with |score| > 0.25.

For *FigR* (v0.1.0)^16^, CRE-Gene interactions were inferred by computing the Spearman’s correlation between the log-normalized gene expression and genomic neighboring CREs accessibility counts (window size = 500k, FDR < 0.1, number of CREs per gene ≥ 2, hg19 RefSeq TSS annotation). Domains of regulatory chromatin (DORCs) scores were computed per gene by summing the accessibility counts of its associated CREs. Both DORC scores and log-normalized gene expression were smoothed using KNN imputation, leveraging SnapATAC2’s spectral embeddings (k = 10). For each DORC, the method identified its k nearest neighbors based on the smoothed DORC scores and pooled their associated peaks. It then calculated TF binding frequencies for these pooled peaks with the MOODs^52,53^ motif matcher algorithm and the CIS-BP^49^ motif database, which were compared to a GC-corrected permuted background set (n = 50) using a z-test to assess significance. For the normalized gene expression, the Spearman’s correlation was calculated between TF expression and DORC accessibility scores. A final regulation score was determined by combining the significance values of the TF–DORC correlation and TF binding prediction, with activation or inhibition assigned based on the sign of the correlation.

For *GRaNIE* (v1.9.7)^17^, pseudo-bulk profiles at the cell-type level were generated for both omics using decoupler-py^54^. Genes were normalized with the quantile normalization from limma^55^, and CREs with the size factors from DESeq2^56^. TF-CRE links were extracted by first overlapping predicted TF binding regions from HOCOMOCOv3^51^ with PWMscan^57^ to all CREs and then computing Pearson’s correlation between normalized gene expression and chromatin accessibility. Similarly, CRE-Gene links were inferred by first assigning CREs to neighboring genes based on genomic distance and computing the Pearson correlation between pairs (window size = 500k). Finally, the two types of connections were merged and filtered for significance (TF-CRE FDR < 0.2, CRE-Gene < 0.2).

For *Pando* (v1.0.5)^18^, CREs were redefined to only include base pairs present in evolutionary conserved regions from phastCons^58^ and not present in annotated exonic regions. TF were assigned to CREs using TF motifs from JASPAR^59^ and CIS-BP^49^ with MOODs^52,53^. CREs were assigned to neighboring genes with the GREAT algorithm^60^ (window size = 500kb). Then, multivariate generalized linear models were fitted for every gene explained by the interaction of the accessibility of an assigned CRE with the expression of a TF assigned to that CRE. This was only done for features with enough TF-CRE or Gene-CRE Pearson’s correlations (|Pearson| > 0.05). Finally, coefficients were filtered for significance, summed to the TF-Gene level and filtered by minimum number of targets (FDR < 0.1, R2 > 0.05, module size > 3).

For *SCENIC+* (v1.0a2)^19^, accessibility count matrices were imputed using pyCisTopic by inferring topics (number of topics = 50)^19^. CREs were assigned to neighbouring genes (window size = 500kb) by predicting observed gene expression by CRE accessibility using GRNBoost2^61^. TF binding was inferred using the CisTarget and DEM motif matcher algorithms, along with the CisTargetDB motif collection^19^. Potential TF-gene links were identified by predicting gene expression from TF expression with GRNBoost2. These links were ranked using a triplet score that incorporated the ranks of CRE-gene, TF-CRE, and TF-gene associations. A final TF-gene score was then computed as (1 - quantile(triplet score)) * sign(regulation).

### Stability score

For features and cells, a stability score was computed for the TFs, edges, and gene overlaps coefficients across the decreasing downsampling iterations. This score was generated for each method by calculating the area under the overlap coefficient curve (AUC). The AUC was determined using the trapezoidal rule applied to a normalized sequence of downsampling steps ranging from 0 to 1, with five steps, and the corresponding overlap coefficient values for that method. This approach provides a quantitative measure of the GRN method’s stability when downsampled by the number of cells or features.

### Unimodal GRN inference methods

This section describes the GRN methods that do not directly model multimodal data, but still can incorporate CRE information through prior knowledge.

Literature-derived GRNs, *CollecTRI*^9^ and *DoRothEA*^8^, were obtained from their Zenodo^62^ and GitHub repositories, respectively. For each dataset, non-expressed genes were removed from the GRNs. Then, a CRE was added for each gene comprising its promoter region (± 1kb from the TSS defined by Ensembl 111^24^). Finally, TF regulons were retained only if they contained more than five target genes.

*SCENIC*^20^ first classified genes as TFs or non-TFs using a custom annotation provided by the method. For each gene, it then fitted an XGBoost model to predict its observed expression based on the expression of TF genes. Next, target genes within each TF regulon were pruned based on TF motif binding enrichment at their promoter regions. Finally, interactions were filtered by score, retaining only those with a score > 0.001.

For each dataset, a random GRN was generated by sampling 25% of the expressed genes at random. For each sampled gene, neighboring accessible CREs were selected within a 500 kb window based on the TSS annotation from Ensembl 111^24^, using an exponential distribution (scale = 1). For each sampled CRE, annotated TFs from Lambert’s database^63^ that were also present in the sampled genes were similarly sampled using an exponential distribution (scale = 1). The resulting elements were merged, and TF regulons were randomly subsampled to achieve a TF-to-gene ratio of 10%.

### Modularized multimodal GRN inference methods

For each original method, we implemented a modularized version designed to be compatible with other inference steps. While these individual steps largely mirrored the original implementations, minor modifications were made in some of the approaches to ensure maximum compatibility across different combinations of methods.

For *CellOracle*, accessibility imputation based on proximities in UMAP coordinates was not performed during CRE-CRE interaction inference with Cicero, as distances in UMAP space have no meaning^64^.

For *Dictys*, datasets preprocessed using alternative strategies also incorporated *Dictys*’ approach to reduce potential sparsity. Furthermore, when building the scaffold GRN prior to modeling, only peaks identified in the CRE-gene association step were included.

For *GRaNIE*, CRE-to-gene associations were performed first, followed by TF binding predictions within those associated CREs, rather than all available CREs. This approach was adopted because many peaks with TF binding predictions in the original implementation were ultimately excluded during the CRE-to-gene merging step. Additionally, the modularized pipeline required a sequential processing order.

Similarly, for *Pando*, the processing order was adjusted from the original implementation: CRE-gene associations were performed first, followed by the TF binding step. Additionally, to ensure compatibility with *FigR*, negative -log(p-values) obtained from MOODS were set to zero in the rare instances where they occurred.

For *SCENIC+*, CRE-gene interactions were filtered based on importance (importance > 2.2e-16). In the TF binding step, only CREs identified in the previous CRE-gene step were considered, rather than the entire set of CREs.

### Infrastructure

All Python-based methods were executed on a high-performance computing cluster using 32 cores, with a maximum memory allocation of 128 GB per step. R-based methods were restricted to a single core, as parallelization was observed to be slower and caused excessive memory consumption. To optimize runtime, individual steps in the modularized pipeline were allowed a maximum execution time of nine hours; steps exceeding this limit returned an empty GRN. Additionally, combinations of steps that failed to produce valid CRE-gene associations, TF binding predictions, or TF-gene predictions also resulted in empty GRNs.

### Evaluation metrics

For the mechanistic metrics, log fold changes and their associated metadata from single-TF perturbations were retrieved from KnockTF^65,66^. For the “TF activity” metric, perturbation experiments were filtered based on terms related to the dataset being evaluated (see https://github.com/saezlab/greta/blob/main/config/prior_cats.json). If a perturbed TF was present in the GRN, TF enrichment scores were inferred using decoupler-py’s ULM method^54^. A perturbation was considered a true positive (TP) if the obtained enrichment score was both positive and significant (BH FDR < 0.05). For the “Forecasting” metric, assays were similarly filtered by matching terms. When a TF was present in the GRN, the effect of the perturbation was simulated by first averaging gene expression values across all cells from the original single-nuclei multiome dataset and then performing in silico perturbation simulations using CellOracle’s perturbation framework (number of steps = 3). Perturbations were considered TPs when the Spearman’s correlation between the simulated changes and the measured changes was significant (ρ > 0.05; BH FDR < 0.05). For the “Steady states” metric, marker TFs were first identified by differential expression analysis at the single-cell level with scanpy^43^ (two-sided Wilcoxon test, BH FDR < 2.22e-16, log2FC > 2) based on the annotated TFs from Lambert^63^. Then, each GRN was filtered to contain only the interactions between these marker TFs. The resulting network was transformed into a set of Boolean rules where we assumed that the activation of a TF depends on the presence of any positive interaction and the absence of every possible inhibition. In more detail, we establish the following formula for a given *TF_A_*: *TF_A_* ∼ (*activator_1_* | *…* | *activator_n_*) & (!*inhibitor_1_* & … & !*inhibitor_m_*), where “|” denoted the operator “OR”, “&” denoted the operator “AND”, and “!” denoted the negative operator. Here, *activators* were TF with positive interaction weights to *TF_A_*, and *inhibitors* were TFs with negative interaction weights to *TF_A_.* For each *TF*, its top 10 regulators based on the absolute value of their interaction weight were considered. With the collection of Boolean rules, all possible steady states were inferred using PyBoolNet^67^(max_output=100k). We then evaluated the overlap of the active TFs in each steady state with the marker TFs of each cell type using a one-sided Fisher’s exact test (BH FDR < 0.01). A steady state was considered a TP when it mapped to only one of the original cell types.

For the predictive metric “Omics,” cells from the original single-nuclei multiome dataset were stratified by cell type, sampled, and split into training and testing sets (train size = 66%, test size = 33%). XGBoost was then used to model the following relationships: Genes ∼ TFs, Genes ∼ CREs, and CREs ∼ TFs. Log-normalized gene expression was used for genes and TFs, while log-normalized chromatin accessibility was used for CREs. After fitting the models, predictions were made on the test set. Features were considered true positives (TPs) if the Spearman correlation was significant (ρ > 0.05; FDR < 0.05).

For the predictive metric “Gene sets,” several collections of gene sets were compiled, including Hallmarks^68^, Kegg^69^, Reactome^70^ and PROGENy^71^. To identify relevant gene sets in the original single-nuclei multiome dataset, gene set enrichment scores were inferred at the cell level using decoupler-py’s ULM method ^54^ and filtered for significance (minimum proportion of cells = 20%; enrichment score > 0; BH FDR < 0.01). For each TF in the GRN, a one-sided Fisher’s exact test was performed to determine which gene sets were overrepresented among its targets (BH FDR < 0.01). A gene set was considered a TP if it was overrepresented in at least one TF regulon within the GRN and was also significant in the original dataset.

For the literature-derived “TF marker” metrics, data from the Human Protein Atlas (HPA)^72^ and TF-Marker^73^ databases were retrieved. Entries from HPA were retained if the protein was annotated as a TF in Lambert^63^, had evidence at the protein level, contained the word “Nucle” in their subcellular location, and was not non-specifically expressed across tissues, cell lines, or single-cell clusters. Both databases were further filtered based on terms relevant to the dataset being evaluated. A TF was considered a true positive (TP) if it was present in both the GRN and the database.

For the literature-derived “TF-TF pair” metric, protein-protein interactions were extracted from EuropePMC^74^ and IntAct^75^. For EuropePMC, the number of times each pair of TFs annotated in Lambert^63^ appeared together, separately, or not at all in available publication titles and abstracts was retrieved using EuropePMC’s RESTful API. This information was then used to compute a one-sided Fisher exact test. Pairs with significant overlap in publications were annotated as interacting pairs (BH FDR < 2.2e-16, Odds ratio >5). For IntAct, interactions were retained if both proteins were annotated as TFs in Lambert^63^ and had a high confidence score (MIscore > 0.75). For a given GRN, TF-TF pairs were identified by testing whether their target genes were overrepresented with respect to each other using a one-sided Fisher’s exact test (BH FDR < 0.01). A TF-TF pair was considered a true positive (TP) if it was present in both the GRN and the database.

For the literature-derived “TF binding” metric, we processed three large databases of ChIP-seq measurements: ChIP-Atlas^76^, Remap2022^77^ and Unibind^78^. From ChIP-Atlas, significant ChIP-seq TF binding peaks were retrieved for the TFs annotated in Lambert^63^(BH FDR < 1e-50; maximum peak length < 750). From Remap2022, the non-redundant ChIP-seq TF binding peaks were retrieved for the TFs annotated in Lambert^63^(maximum peak length < 750). From UniBind, ChIP-seq TF binding peaks from the robust collection were retrieved for the TFs annotated in Lambert^63^(maximum peak length < 750). For each database, overlapping peaks from the same TF were merged using bedtools^79^. For a given dataset and GRN, TF binding databases were filtered to include only ChIP-seq peaks overlapping measured CREs from the original chromatin accessibility data using pyranges (v0.1.2)^80^, TFs expressed in the original gene expression data, and cell type or tissue terms relevant to the dataset being evaluated. A TF binding peak was considered a TP if it overlapped with a CRE regulated by that TF according to the GRN.

For the literature-derived “CREs” metric, a collection of problematic genomic regions (Blacklisted^81^), two enhancer annotation (ENCODE^82^ and Zhang21^83^), an evolutionary conservation (PhastCons^58^) and a SNP-trait (GWAS catalog^84^) databases were processed. Additionally, a promoter resource was generated by opening genomic windows on the TSS annotated by Ensembl 111^24^ (± 1kb from the TSS). For a given dataset and GRN, CRE databases were filtered to include only regions overlapping measured CREs from the original chromatin accessibility data using pyranges (v0.1.2)^80^. A CRE was considered a TP if it overlapped with a CRE in the GRN. For the Blacklisted resource, a CRE was considered a TP if it did not overlap with any of the annotated regions.

For the literature-derived “Genes” metric, eQTL Catalogue^85^ was processed. CRE-Gene interactions were kept if the gene was annotated in Ensembl 111^24^ and the link was significant (FDR < 1e-5). For a given dataset and GRN, the database was filtered to include only genes expressed in the original gene expression data, and cell type or tissue terms relevant to the dataset being evaluated. A CRE-Gene link was considered to be a TP when the interaction was present in both the GRN and the dataset.

### Statistical testing

Statistical tests were performed using scipy (v1.14.1)^86^ and decoupler (v1.8.0)^54^. For false discovery rate (FDR) correction, the Benjamini-Hochberg (BH) procedure was employed.

In the evaluation of the modularized GRN inference runs, to test for significantly ranked approaches the Weighted Kolmogorov-Smirnov-like statistic was repurposed from GSEA^87^ using decoupler’s implementation. In this case, the ranking value was the distribution of F_0.1_ scores across modularized runs for a particular metric-database. The set of interest were the runs containing that specific inference step, for example all runs with TF binding prediction from *Dictys*, and the background were all the runs without that specific step (runs without the TF binding prediction from *Dictys*). By computing the Weighted Kolmogorov-Smirnov-like statistic, an enrichment score was obtained which was tested using random permutations of background runs (number of permutations = 1000). Significance was assigned based on the sign of the enrichment score (score > 0; BH FDR < 0.01).

### Plots and figures

Plots were generated using the matplotlib (v3.9.2)^88^, seaborn (v0.13.2)^89^ and marsilea(v0.3.2)^90^ libraries. Figures were then edited with Inkscape.

## 5. Acknowledgements

Thanks to Christian Arnold, Seppe de Winter, Adam Kyle, Philipp Sven Lars Schäfer, Aryan Kamal, Jalil Nourisa, Manu Saraswat, Anshul Kundaje, Judith Zaugg, Stein Aerts, Carl Herrmann, Yvan Saeys, and Ingrid Lohmann for helpful discussions about gene regulatory networks. We also thank the method developers who supported us with the technical implementation of their tools.

SMD was supported by the German Federal Ministry of Education and Research, particularly the LiSyM Cancer research core (BMBF, Funding number: 031L0257B). The authors acknowledge support by the state of Baden-Württemberg through bwHPC and the German Research Foundation (DFG) through grant INST 35/1597-1 FUGG. They also gratefully acknowledge the data storage service SDS@hd supported by the Ministry of Science, Research and the Arts Baden-Württemberg (MWK) and DFG through grant INST 35/1503-1 FUGG.

## 7. Authors contributions

PBM, RRF and JSR conceived and designed the study. PBM, LW and SMD searched and processed the single-cell multiome datasets. PBM, RCF, LW and RRF searched and processed the evaluation databases into metrics. PBM, RCF, RT and YY implemented the multimodal gene regulatory inference methods. PBM performed the computational analyses, supervised by RRF and JSR. PBM, RRF and JSR wrote the manuscript. PBM and SMD illustrated and organized the figures. All authors have read, edited and approved submission of the manuscript.

## Supplementary Materials

## Supplementary note 1

The downsampling strategy also enabled the quantification of the computational cost of running these methods (**Extended Data Fig. 2d**). Excluding the literature-derived GRNs and the random baseline, the majority of methods took less than three hours to run on 32 CPUs (full dataset mean running time = 3.613 hours), the exceptions being *Dictys* which also required a GPU, and *FigR*, which we found that its computing time grew exponentially with the number of cells and features (full dataset mean running time: *Dictys* = 6.84 hours, *FigR* = 12.69 hours). Regarding memory usage, most methods required more than 16 GBs (full dataset mean memory = 19.765 GBs), the exceptions being *Dictys* and *GRaNIE* (full dataset mean memory: *Dictys* = 11.63 GBs, *GRaNIE =* 4.6 GBs).

---

**Extended Data Table 1.**
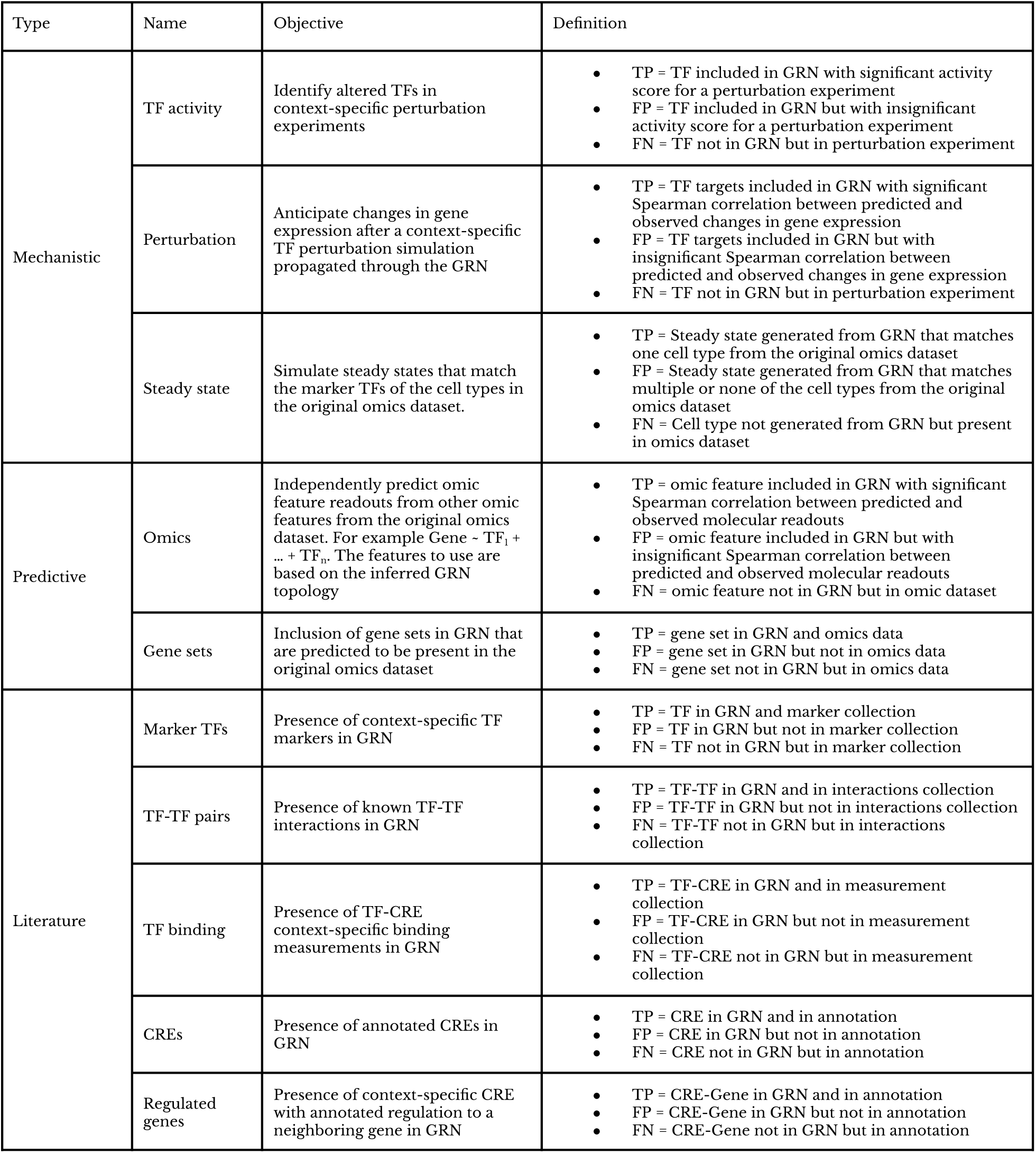
Definition of evaluation metrics. List of evaluation metrics used to assess the performance of GRNs together with their objective and definition. Metrics that mention context-specific information were tailored for each specific dataset. For example, when working with a heart dataset only entries that contain cardiac-related labels would be used. True Positives (TP), False Positives (FP) and False Negatives (FN).

| Type | Name | Objective | Definition |
| --- | --- | --- | --- |
| Mechanistic | TF activity | Identify altered TFs in context-specific perturbation experiments | TP = TF included in GRN with significant activity score for a perturbation experiment<br>FP = TF included in GRN but with insignificant activity score for a perturbation experiment<br>FN = TF not in GRN but in perturbation experiment |
|  | Perturbation | Anticipate changes in gene expression after a context-specific TF perturbation simulation propagated through the GRN | TP = TF targets included in GRN with significant Spearman correlation between predicted and observed changes in gene expression<br>FP = TF targets included in GRN but with insignificant Spearman correlation between predicted and observed changes in gene expression<br>FN = TF not in GRN but in perturbation experiment |
|  | Steady state | Simulate steady states that match the marker TFs of the cell types in the original omics dataset. | TP = Steady state generated from GRN that matches one cell type from the original omics dataset<br>FP = Steady state generated from GRN that matches multiple or none of the cell types from the original omics dataset<br>FN = Cell type not generated from GRN but present in omics dataset |
| Predictive | Omics | Independently predict omic feature readouts from other omic features from the original omics dataset. For example Gene ~ $TF_1 + \dots + TF_n$. The features to use are based on the inferred GRN topology | TP = omic feature included in GRN with significant Spearman correlation between predicted and observed molecular readouts<br>FP = omic feature included in GRN but with insignificant Spearman correlation between predicted and observed molecular readouts<br>FN = omic feature not in GRN but in omic dataset |
|  | Gene sets | Inclusion of gene sets in GRN that are predicted to be present in the original omics dataset | TP = gene set in GRN and omics data<br>FP = gene set in GRN but not in omics data<br>FN = gene set not in GRN but in omics data |
| Literature | Marker TFs | Presence of context-specific TF markers in GRN | TP = TF in GRN and marker collection<br>FP = TF in GRN but not in marker collection<br>FN = TF not in GRN but in marker collection |
|  | TF-TF pairs | Presence of known TF-TF interactions in GRN | TP = TF-TF in GRN and in interactions collection<br>FP = TF-TF in GRN but not in interactions collection<br>FN = TF-TF not in GRN but in interactions collection |
|  | TF binding | Presence of TF-CRE context-specific binding measurements in GRN | TP = TF-CRE in GRN and in measurement collection<br>FP = TF-CRE in GRN but not in measurement collection<br>FN = TF-CRE not in GRN but in measurement collection |
|  | CREs | Presence of annotated CREs in GRN | TP = CRE in GRN and in annotation<br>FP = CRE in GRN but not in annotation<br>FN = CRE not in GRN but in annotation |
|  | Regulated genes | Presence of context-specific CRE with annotated regulation to a neighboring gene in GRN | TP = CRE-Gene in GRN and in annotation<br>FP = CRE-Gene in GRN but not in annotation<br>FN = CRE-Gene not in GRN but in annotation |

**Extended Data Figure 1.**
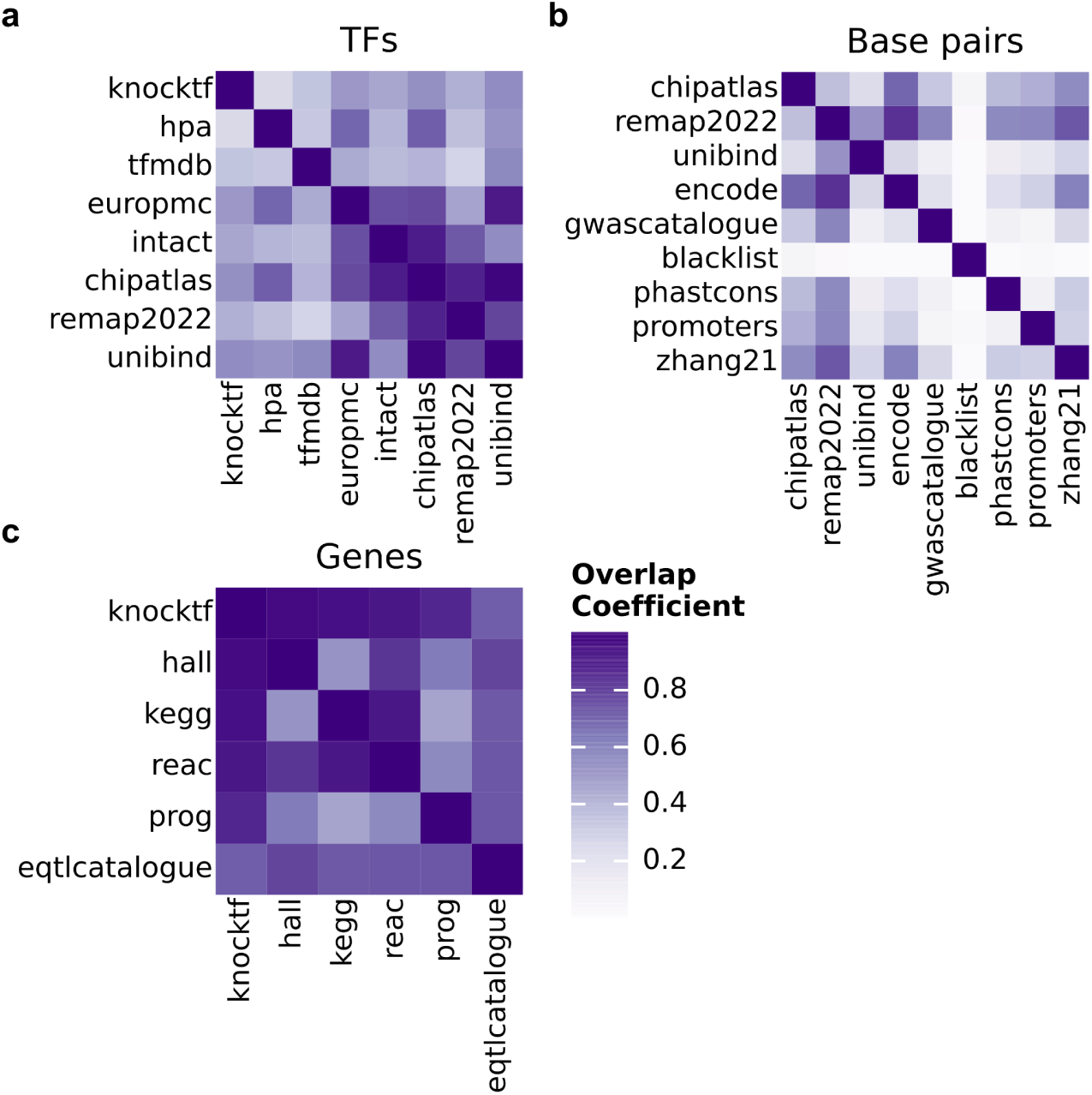
Cover age across evaluation databases. Overlap coefficient of TFs (**a**), base pairs (**b**) and target genes (**c**) across the evaluation databases.

**Extended Data Figure 2.**
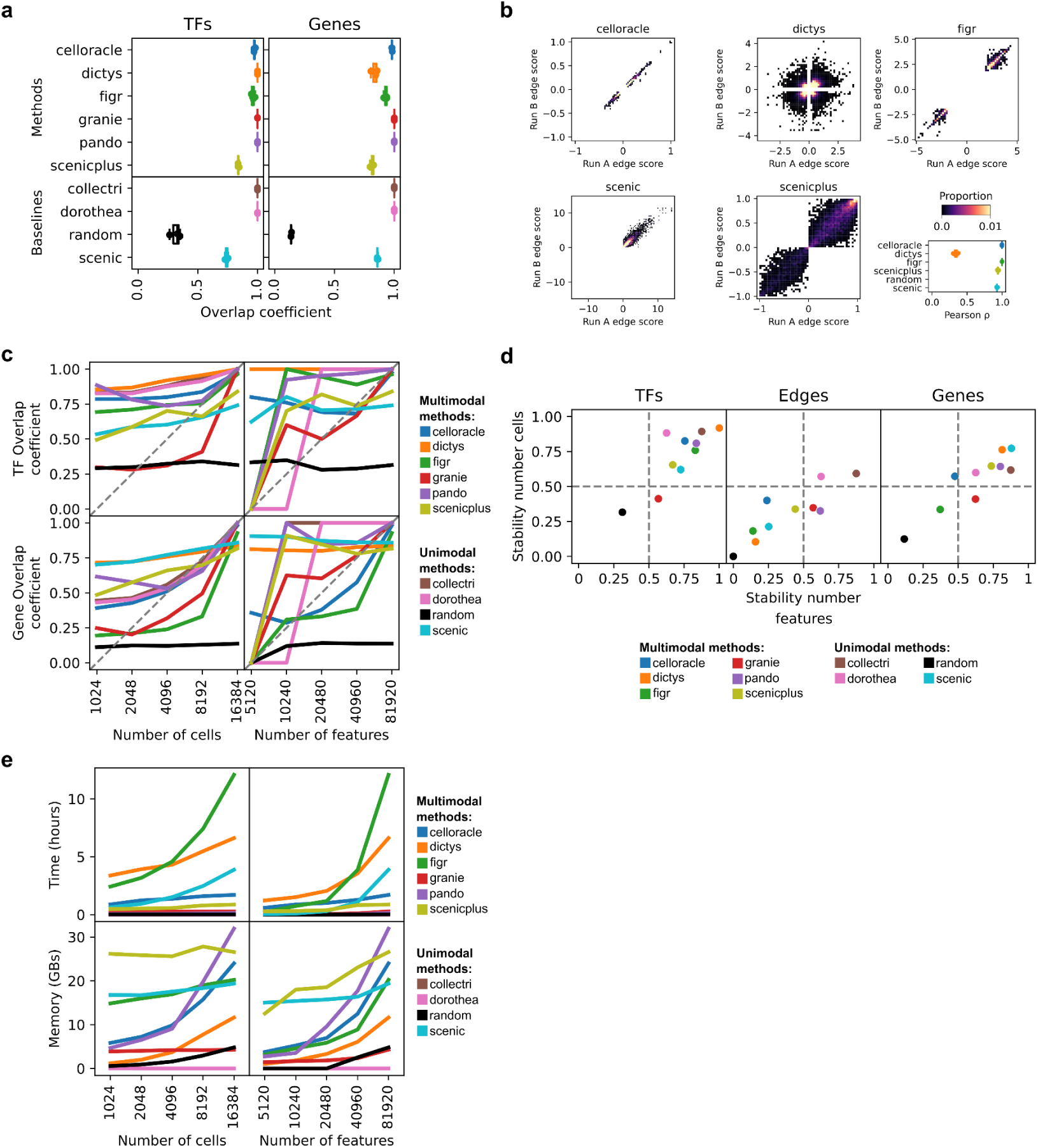
**a,** Overlap coefficients of TFs (left) and target genes (right) between three separate runs per method. **b,** Example comparison of edge interaction scores for the non-deterministic methods when using the same dataset (left) and the Pearson ρ between the three different runs (right). **b,** Overlap coefficients of TFs (top) and target genes (bottom) between GRNs inferred from downsampled datasets with the one inferred using all available data. **c,** Stability scores for TFs, edges and target genes. The x-axis indicates the stability score when the number of cells is fixed and features are downsampled, while the y-axis indicates the stability score when the number of features are fixed and cells are downsampled. **e,** Mean time (top) and memory (bottom) used across methods for each downsampled dataset.

**Extended Data Figure 3.**
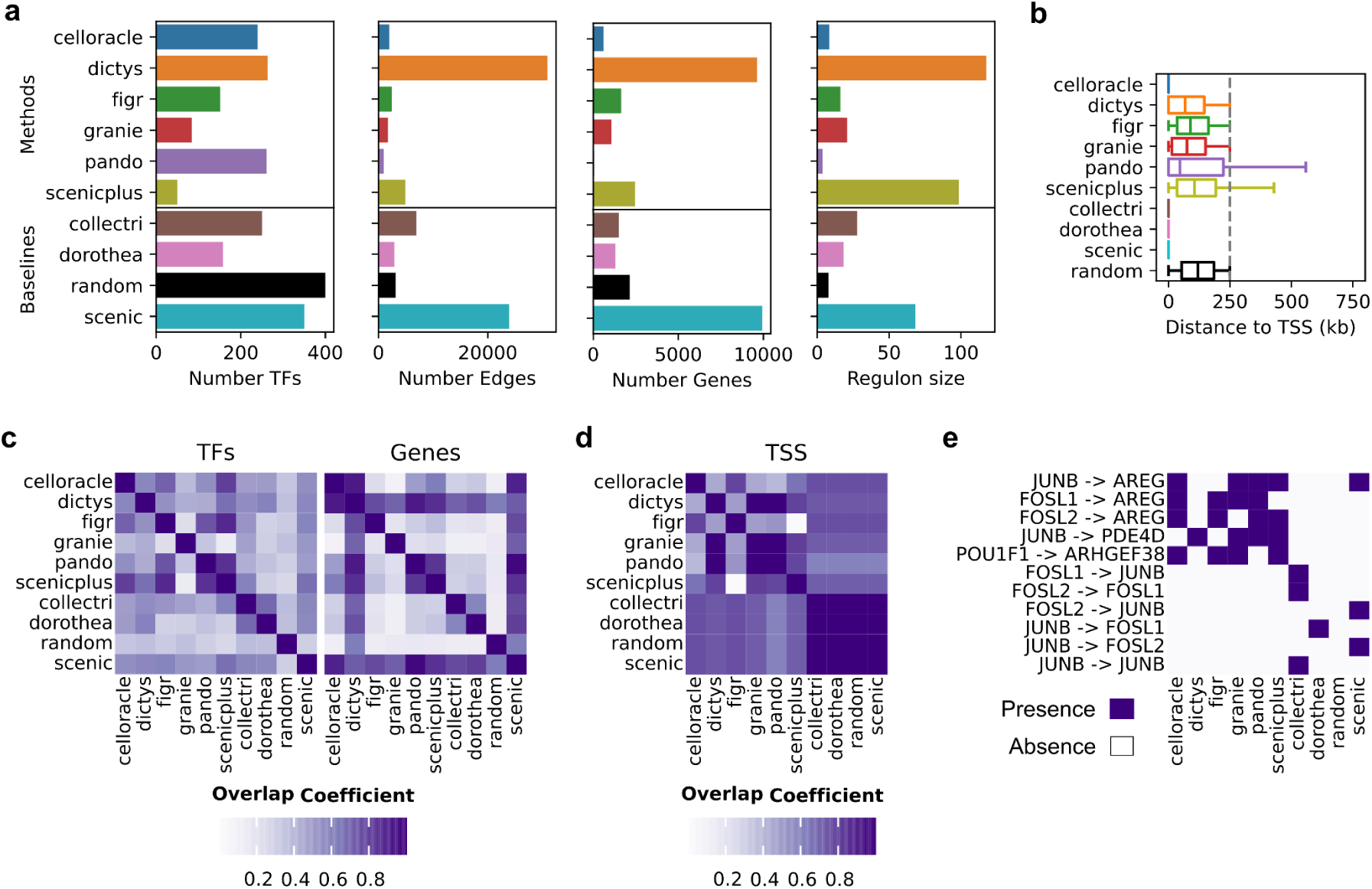
**a,** Number of transcription factors (TFs) (far left), number of edges (left), number of genes (right), and average number of target genes per TF (far right) across methods. **b,** Distances of Cis-regulatory elements (CRE) to transcription starting sites (TSS) by method. Dashed line indicates the window size used in all methods (± 250 kb around the TSS). **c,** Overlap coefficients of TFs (left) and genes (right) across methods. **d,** Genomic overlap coefficient of annotated TSS regions across methods. **e,** Binary adjacency matrix of interactions shared across at least three multimodal methods.

**Extended Data Figure 4.**
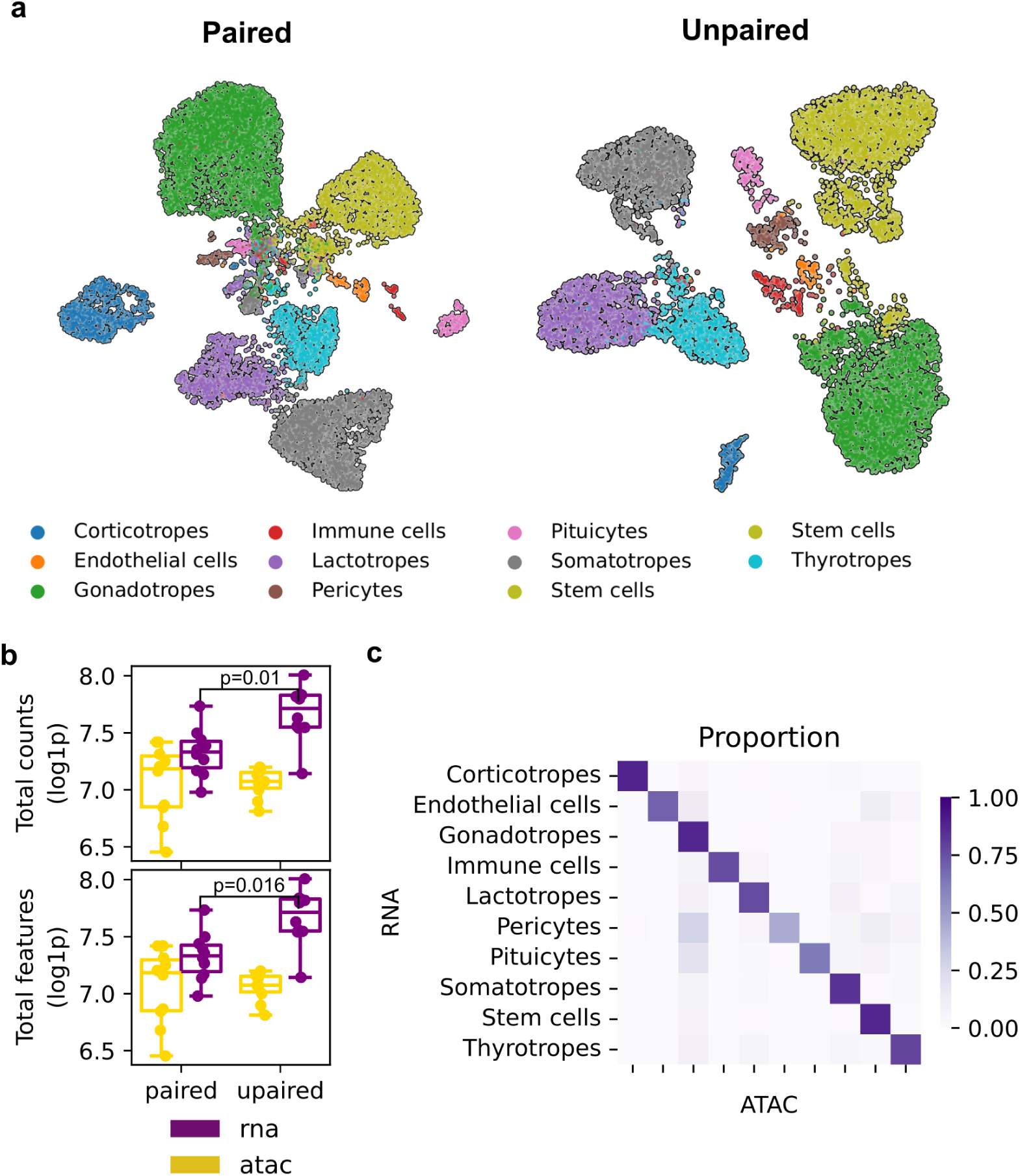
Similarities between paired, unpaired and synthetic pituitary datasets. **a,** UMAP visualization of the cell type clusters for the paired (left) and unpaired (right) datasets. Color represents the different cell types. **b,** Mean total number of counts and mean number of features with counts per cell type for the paired and unpaired dataset. **c,** Proportion of barcodes from one cell type matched to the others after integration in the synthetic paired dataset.

**Extended Data Figure 5.**
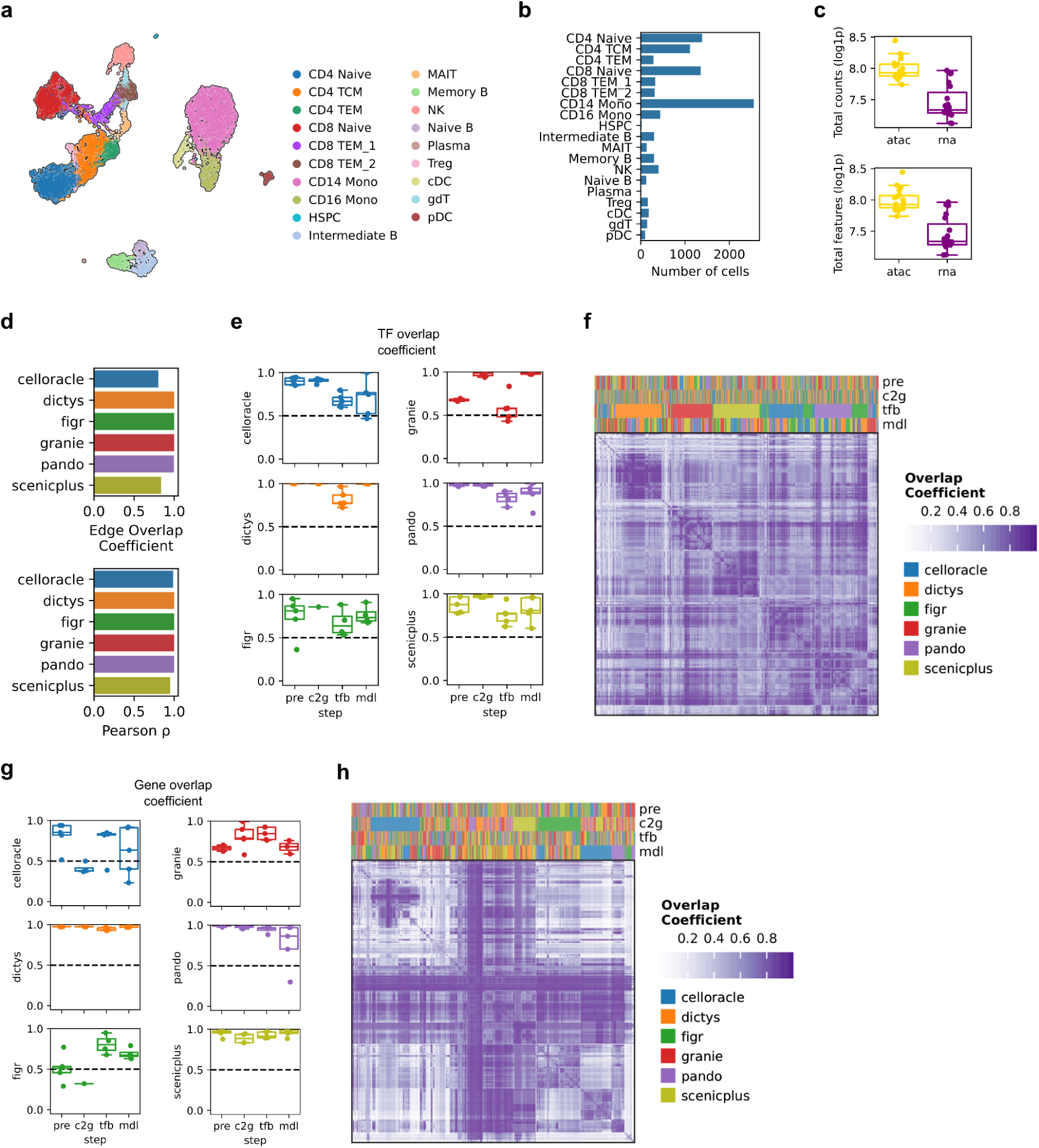
Peripheral blood mononuclear cell dataset and similarities between runs. **a,** UMAP visualization of the cell type clusters for the peripheral blood mononuclear cell dataset. Color represents the different cell types. **b,** Number of cells per cell type. **c,** Mean total number of counts and mean number of features with counts per cell type. **d,** Overlap coefficient at the edge level (top) and Spearman’s ρ (bottom) between the original methods and their equivalent modularized counterpart (top). **e,** Effect of changing one of the inference steps from a fixed method pipeline to another method, measured by the overlap coefficient at the TF level. **f,** Pairwise overlap coefficient at the TF level between all possible inference runs combinations. **g, h** are the same as e and f but for the overlap coefficient at the gene level.

**Extended Data Figure 6.**
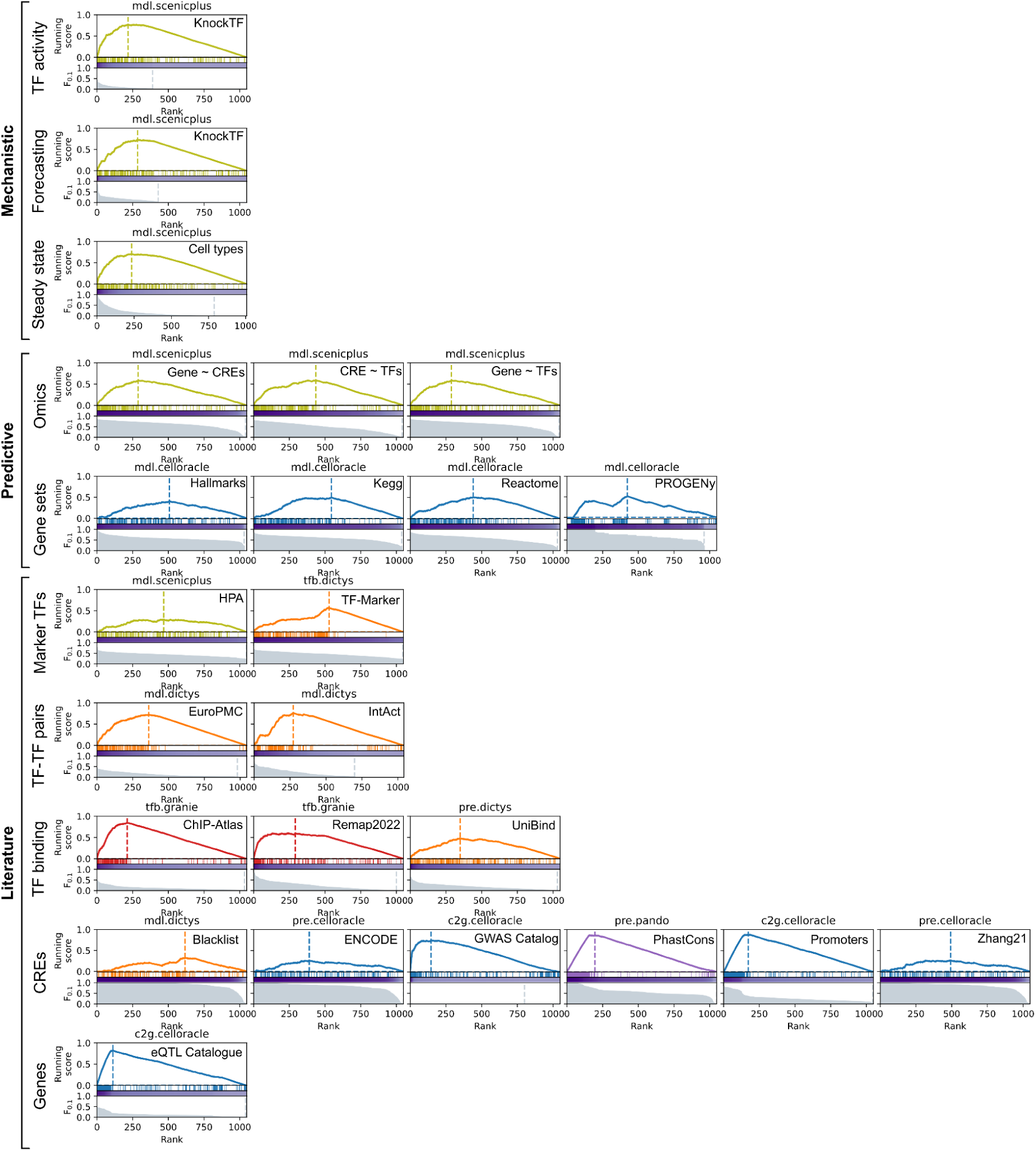
Weighted Kolmogorov-Smirnov-like statistics plots. Weighted Kolmogorov-Smirnov-like statistic plots of the inference step method combination with the most significant p-value for each evaluation task.

**Extended Data Figure 7.**
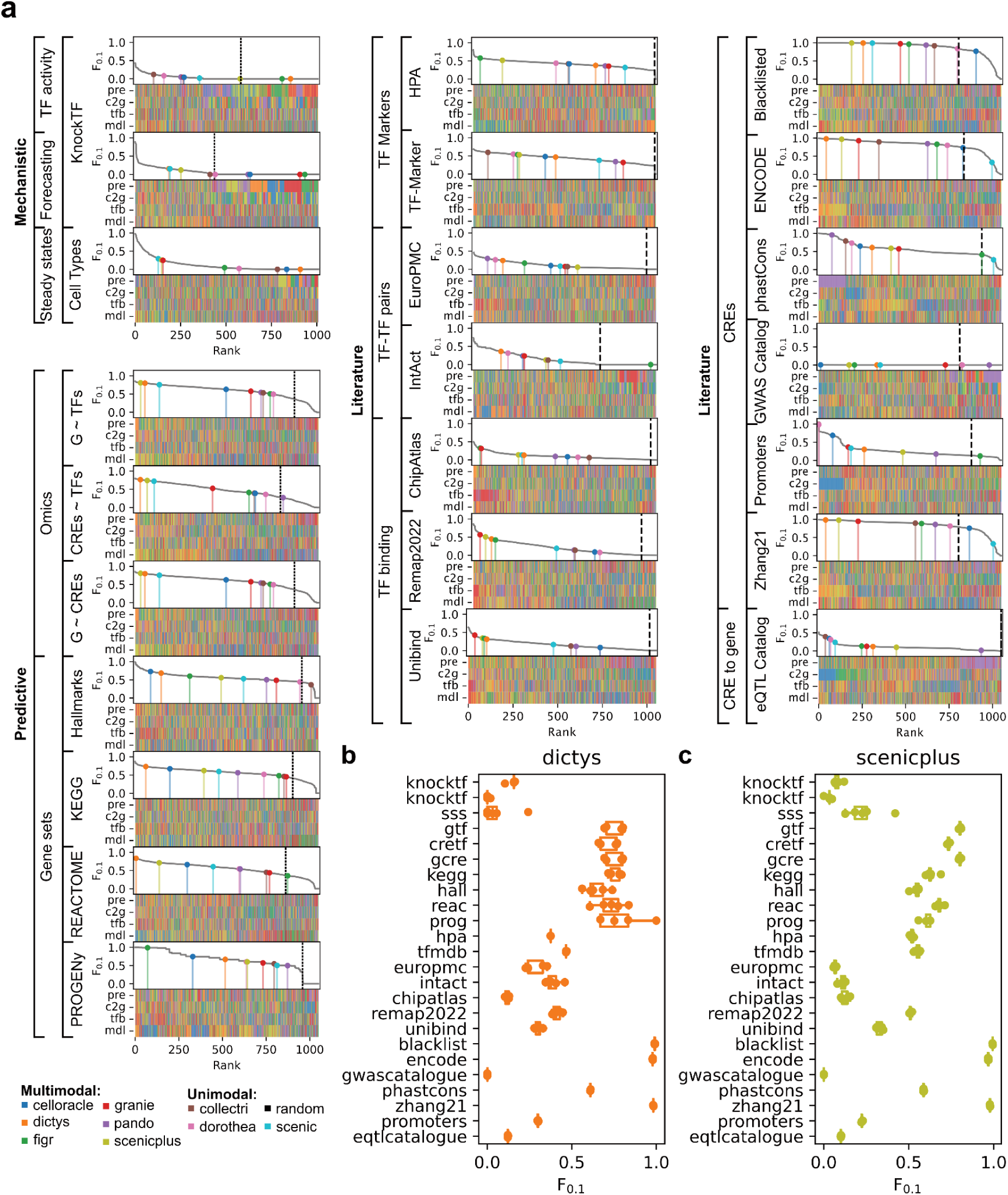
Performance of fixed multimodal and unimodal methods. **a,** Ranks of fixed multimodal and unimodal methods across evaluation tasks. **b,c,** Stability of performances for fixed Dictys runs and SCENIC+ using five different random seeds.

